# Circular RNA profiling revealed an evolutionarily conserved circACACA promotes liver lipid metabolism, oxidative stress, and autophagy disorder in a ceRNA manner

**DOI:** 10.1101/2025.05.15.654219

**Authors:** Jing Zhao, Shunshun Han, Jialin Xiang, Yuqi Chen, Xiyu Zhao, Wenjuan Wang, Yao Zhang, Qing Zhu, Chang Liu, Huadong Yin

## Abstract

Non-alcoholic fatty liver disease (NAFLD) is a clinical syndrome characterized primarily by hepatocellular steatosis and lipid accumulation, which leads to hepatocyte apoptosis, autophagy, inflammation, and intracellular oxidative stress. NAFLD is recognized as one of the most prevalent and complex chronic liver diseases globally, with its occurrence and associated mortality rates rising swiftly each year. Due to the high similarity between chicken fatty liver syndrome (FLS) and NAFLD, as well as the easy availability of diseased chickens, the chicken is considered an ideal model for studying the pathogenesis of NAFLD. Previous studies have pinpointed several circular RNAs (circRNAs) implicated in the pathogenesis of NAFLD, yet the underlying functions and mechanisms of numerous circRNAs continue to remain elusive. In this experiment, we utilized circRNA sequencing of chicken livers to identify a novel circRNA, named circACACA, and discovered that it disrupts the metabolic homeostasis of lipids within hepatocytes. Consequently, this disruption leads to oxidative stress and the induction of autophagy, ultimately exerting an adverse effect on chicken liver health. Mechanistically, circACACA functions as a molecular sponge for miR-132b-5p and miR-101-2-5p to modulate the expression of the downstream CBFB/PIM1 complex. Consequently, it influenced the activity of the AKT/mTOR and PPAR-γ signaling pathways to perform its physiological functions. Crucially, we noticed substantial sequence similarity of circACACA across diverse species by comprehensively searching databases. Further, our research with a mouse model confirmed that the functional conservation of circACACA across livers of different species. Overall, this study built a mechanistic network for circACACA and confirmed its sequence conservation and functional relevance across various species. Our results not only provide new targets for the prevention and treatment of NAFLD but also present fresh perspectives for progress in healthy production of laying hens.

## Introduction

Non-alcoholic fatty liver disease (NAFLD) currently stands as the primary cause of chronic liver diseases worldwide, impacting 25% of the global populace and roughly 70 to 80% of individuals who are obese or diabetic [1, 2]. Diagnosis of NAFLD occurs when fat accumulation in the liver exceeds 5% to 10% of hepatocytes, in the early stages, most individuals with NAFLD are asymptomatic, with hepatic steatosis that typically does not evolve into a more severe disease [3]. Nevertheless, as excessive fat accumulates in the liver, which triggers oxidative stress leading to further exacerbation of liver cell damage and inflammatory responses, known as non-alcoholic steatohepatitis (NASH), may potentially deteriorate into advanced liver diseases, including cirrhosis or even hepatocellular carcinoma (HCC) [4–6]. This complex pathological process is triggered by various factors including environmental, genetic and nutritional influences. Current management and prevention strategies primarily focus on reducing caloric intake and enhancing physical activity [7]; However, these measures do not address underlying causes of the disease. Numerous studies have revealed the regulatory role of genetic factors in susceptibility to NAFLD indicating their crucial contribution to development of this condition.

In chickens, a condition analogous to NAFLD is termed fatty liver syndrome (FLS) [8]. Both FLS and NAFLD exhibit similar pathogenic mechanisms, stemming from excessive lipid accumulation in the liver, ultimately causing hepatic steatosis and metabolic dysfunction [2, 9]. Furthermore, both conditions display pathological characteristics such as hepatocyte swelling, distortion, and fibrosis [10, 11]. Notably, chickens can readily induce FLS through high-fat diets or feed restriction, offering a controllable and consistent model for investigating the onset and progression of NAFLD [12, 13]. In comparison to large mammals, chickens offer cost-effectiveness and ethical advantages, rendering them highly suitable for extensive research into metabolic diseases [11]. Therefore, the chicken is considered an excellent biological model for studying NAFLD.

Circular RNAs (circRNAs) represent a distinct class of noncoding RNAs characterized by their closed and stable loop structures formed via back splicing events [14]. Owing to their remarkable conservation, inherent stability, and tissue-specific expression patterns, circRNAs hold great promise as diagnostic biomarkers and therapeutic targets for various diseases, including NAFLD [15, 16]. Increasing evidence indicates that abnormal expression of circRNAs is associated with the emergence and progression of liver diseases, encompassing hepatic steatosis, liver regeneration and hepatocellular carcinoma [16–18]. In recent years, three common NAFLD models including high-fat diet models, methionine, choline-deficient (MCD) models and HepG2 cell line model, have been used to study NAFLD, and several circRNAs have been found to be differentially expressed between the model and control group [19–21]. However, the circRNAs with definitive regulatory functions and their underlying mechanisms in NAFLD still need further identification. Given the high level of conservation observed among the majority of circRNA sequences, prioritizing those circRNAs that are conserved across species within animal sequence profiles could pave the way for novel advancements in clinical trials for humans.

In this study, we initially conducted the circRNA sequencing on a chicken model with FLS to identify candidate circRNAs that regulate this condition in chickens.

Subsequently, the screened circRNAs were functionally validated in chicken hepatocytes. Finally, a mouse model was utilized to verify the interspecies conservation of the selected circRNAs in regulating liver metabolism. This study can provide new strategies for investigating the molecular mechanisms of circRNA regulation in NAFLD.

## Materials and methods

### Animal and sample collection

The experimental protocols in this research were approved by the Animal Welfare Committee of Sichuan Agricultural University, with the approval number 2021102006. The experiment utilized a total of one hundred 200-day-old Tianfu layers provided by the poultry breeding farm at Sichuan Agricultural University (Ya’an, Sichuan, China). As our previous reported, these chickens were randomly divided into two groups, the chickens in the experimental group were allowed to freely consume feed according to their nutritional requirements to establish a fatty liver (FL) model, whereas those in the control group received a daily feed allowance of 120 g to obtain the control liver (CL) [22]. When the chickens reached 300 days of age, three individuals were randomly chosen from each group for RNA extraction and circRNA sequencing. Tissues including the liver, lung, brain, spleen, kidney, abdominal fat, breast muscle, leg muscle, and intestine were excised in the control group and frozen at -80°C for subsequent analysis.

Sixteen 9-week-old C57BL/6J mice were acquired from DOSSY Experimental Animals (DOSSY, Chengdu, China). And mice were subsequently randomized into two groups. One group was fed a high-fat diet, while the other received a normal diet. Following a three-month treatment period, all mice were euthanized. Blood samples and liver tissues were collected from both groups and preserved for future experiments. Furthermore, RNA extraction was performed on various tissues (liver, spleen, lung, ovary, kidney, breast muscle, leg muscle, brain, intestine, and fat) collected from three mice in the normal diet group.

### CircRNA sequencing

Total RNA extraction from the liver samples was carried out using TRIzol Reagent (Invitrogen, Shanghai, China). Preliminary quantification and assessment of RNA concentration and purity were conducted utilizing a NanoDrop 2000 spectrophotometer. To ensure the quality of the sample library, the insert fragment length of the library was measured using both Qubit and an Agilent 2100 bioanalyzer. Following pooling based on the effective concentration of the library and the data output requirements, Illumina PE150 sequencing was performed. Once the raw sequencing reads were obtained, the quality of the sequencing data was rigorously assessed. Subsequently, bioinformatics analysis was conducted, encompassing quality control, sequence alignment, quantification, and annotation of circRNA. The sequencing data were analyzed with reference to the gallus_gallus_Ensembl_97 genome (ftp://ftp.ensembl.org/pub/release97/fasta/gallus_gallus/dna/).

### CircRNAs identification and differentially expressed analysis

The identification of circRNAs was accomplished through the combined use of Find_circ and CIRI software. The expression levels of both known and novel circRNAs in each sample were statistically analyzed and normalized using Transcripts Per Million (TPM). Differential expression analysis of circRNAs was conducted using DESeq2, which is based on the negative binomial distribution. A significance threshold of an adjusted *P*-value (*P* adj) less than 0.05 was established as the criterion for identifying differentially expressed circRNAs. The TPM values of these differentially expressed circRNAs from all comparison groups were gathered and subjected to hierarchical clustering analysis. Following the identification of differentially expressed circRNAs, Gene Ontology (GO) and Kyoto Encyclopedia of Genes and Genomes (KEGG) enrichment analyses were independently performed, based on the correlation between circRNAs and their respective source genes.

### Histomorphology and Oil red O stain

The liver tissues of chicken and mice liver tissues were fixed using 4% paraformaldehyde (Beyotime, Shanghai, China). Following fixation, dehydration of the samples was achieved through the use of varying concentrations of ethanol. Subsequently, the paraffin-embedded tissue samples were sectioned into 5 um slices and stained with both hematoxylin-eosin (HE) and Oil red O. Random snapshots of the stained images were captured and stored using an electronic microscope.

### Cells culture and transfection

Chicken hepatocytes were isolated from 200-day-old layers following established protocols [22]. Briefly, the intact liver was dissected from the abdominal cavity of the chicken and thoroughly rinsed with calcium-free HEPES buffer (pH 7.5), followed by a calcium-containing washing solution (pH 7.5) until all traces of blood were removed. A selected portion of the liver tissue was then digested using type II collagenase and pancreatin. The hepatocytes were collected by centrifuging the digested tissue at 2000 rpm for 5 min. The cells were cultivated in M199 medium enriched with 10% fetal bovine serum (Gibco, Grand Island, NY, USA) and 1% penicillin-streptomycin (Solarbio, Beijing, China). Optimal cell growth was maintained in a humidified environment at 37℃ with 5% CO_2_, and the medium was replaced every 24 h. Additionally, Alpha Mouse Liver 12 (AML12) cells were cultured in DMEM/F12 medium supplemented with 10% fetal bovine serum, 10 µg/ml insulin, 5.5 µg/ml transferrin, 40 ng/ml dexamethasone, 5 ng/ml selenium, and 1% penicillin-streptomycin (Procell Life Science & Technology, Wuhan, China), under the same culture conditions as those used for the chicken hepatocytes.

For transfection, we utilized Lipofectamine 3000 (Invitrogen) and Opti-MEM® medium (Gibco) to culture the overexpression vector and small interfering RNA (siRNA), adhering to the methodology outlined in a previous study by Zhao et al. [38]. The overexpression vectors were exclusively designed and synthesized by Tsingke (Beijing, China), the siRNAs were synthesized by GenePharma (Shanghai, China). The detailed sequences of these constructs are provided in S1 Table.

### cDNA synthesis and quantitative real-time PCR (qPCR)

Following the extraction of total RNA, cDNA synthesis was carried out using specific kits: the One Step miRNA cDNA Synthesis Kit (HaiGene, Harbin, China) for miRNA and the PrimeScript™ RT Reagent Kit (TaKaRa Biotechnology, Tokyo, Japan) for mRNA. The resulting cDNA served as the template for qPCR testing, following previously reported methods [23]. The qPCR analysis aimed to quantify the mRNA expression levels of the target genes, employing One Step TB Green® PrimeScript™ RT-PCR Kit II (TaKaRa) on the CPX Connect Real-Time System (Bio-Rad, Hercules,

CA, USA). *β-actin*, *GAPDH*, and *U6* were used as internal reference genes, with the detailed primer sequences provided in S2 Table.

### CircRNA verification

To confirm the loop structure of circRNAs, primers specifically designed to span the junction site (termed divergent primers) were synthesized. The cDNA amplified using these divergent primers was then subjected to Sanger sequencing to verify the authenticity of the junction site. Additionally, an RNase R digestion assay was employed to assess the stability of circRNAs. The RNA extracted from liver tissues or hepatocytes was divided into two aliquots. One aliquot was incubated with 1 U/μl RNase R, while the other served as a negative control by being treated with sterile water. Following treatment, all samples were reverse transcribed to generate cDNA, and their expression levels were analyzed using qPCR. Furthermore, RNA samples were reverse transcribed into cDNA using two different methods: one utilizing random N9 primers and the other using oligo d(T) primers. The qPCR was then employed to detect the expression levels of the genes.

### ELISA assay

Chicken ELISA kits sourced from Newgeorge (Shanghai, China) were utilized to measure the activity levels of glutathione (GSH), superoxide dismutase (SOD), total antioxidant capacity (T-AOC), and methane dicarboxylic aldehyde (MDA), following the manufacturer’s instructions. For mouse serum and cellular analysis, biochemical reagent kits provided by AIDISHENG Biological Company (Jiangsu, China) were employed to detect SOD, T-AOC, GSH, and MDA.

### Western blot

The protein concentration extracted from cells was determined using the Bicinchoninic Acid (BCA) protein assay kit provided by BestBio (Shanghai, China). Subsequently, the proteins were separated by sodium dodecyl sulfate-polyacrylamide gel electrophoresis (SDS-PAGE) and transferred onto a polyvinylidene fluoride (PVDF) membrane (Millipore Corporation, Massachusetts, USA). The protein bands were then incubated with specific primary antibodies, followed by the addition of the corresponding secondary antibodies. To enhance the visibility of the protein bands, the Enhanced Chemiluminescence (ECL) reagent from Beyotime was applied. Image capture was facilitated using Image J software to obtain grayscale images. β-tubulin and GAPDH served as internal reference proteins. Detailed information on the antibodies used is provided in S3 Table.

### Co-Immunoprecipitation (Co-IP)

The CBFB-Flag vector was constructed and successfully transfected into cells. Immunoprecipitation was performed according to the protocol outlined in the Pierce™ Co-Immunoprecipitation Kit (Thermo Fisher Scientific, Massachusetts, USA). In brief, cells were harvested, and total proteins were extracted using IP lysis buffer. Subsequently, the IgG antibody was immobilized onto the enhanced AminoLink resin. This was followed by the immunoprecipitation process, elution of the bound proteins, and preparation of SDS-PAGE samples for subsequent western blot analysis.

### Lipid assay

The intracellular content of triglycerides (TG) and total cholesterol (TC) serves as an indicator of intracellular fat accumulation. Commercial biochemistry kits sourced from Jiancheng Bioengineering Institute (Nanjing, China) were used to assess the levels of TG and TC in hepatocytes, adhering strictly to the manufacturer’s instructions. For BODIPY 493/503 staining, cells were plated in a 96-well plate and fixed with 4% paraformaldehyde (Beyotime). Following fixation, the cells were stained with BODIPY 493/503 staining reagent (diluted to 1 µg/ml in PBS; Maokangbio, Shanghai, China) to visualize lipid droplets within the hepatocytes. Hoechst staining was used to stain the cell nuclei. Finally, images of the stained cells were randomly captured using a fluorescence microscope.

### Flow cytometric analysis

Flow cytometry was employed to quantify intracellular reactive oxygen species (ROS) levels. Briefly, hepatocytes were harvested post-transfection and subsequently labeled with the ROS Assay Kit (Beyotime) to detect ROS. The corresponding fluorescence intensities were then measured using a CytExpert flow cytometer. The acquired data were processed and analyzed utilizing Kaluza 2.1 software (Thermo Fisher Scientific).

### Confocal microscope and transmission electron microscopy

Hepatocytes were plated in the 6-well plate and infected with Mcherry-EGFP-LC3 adenovirus. Following the 48 h incubation period, confocal fluorescence microscopy was used to observe the fluorescence activity within the cells. For transmission electron microscopy (TEM) analysis, the transfected cells were harvested after 48 h and fixed using a fixation solution. Subsequently, a JEM-1400 TEM (JEOL, Tokyo, Japan), equipped with an AMT CCD camera (Sony, Tokyo, Japan), was utilized to observe and capture images of the intracellular autophagosomes and lysosomal status.

### Fluorescence in situ hybridization (FISH)

FISH probes targeting the cross-splicing site of circACACA, as well as the sequences of miR-132b-5p and miR-101-2-5p, were designed and synthesized by Tsingke (Beijing, China). The precise sequences of these FISH probes are provided in S4 Table. Cells were plated in dishes containing coverslips and, upon reaching 70-80% confluence, the adherent cells were harvested and fixed. Following fixation, the hepatocytes were labeled with fluorescent probes diluted in DEPC water. The cell nuclei were counterstained with 4’, 6-diamidino-2-phenylindole (DAPI). Subsequently, the desired images were visualized and captured using a confocal fluorescence microscope (Olympus, Melville, NY, USA).

### CircRNA-miRNA-mRNA interaction network construction

The target relationship between circRNA and miRNA was predicted using RNAhybrid. Additionally, miRDB and targetScan were employed to predict the target genes of the miRNAs. The predicted target relationship was further evaluated through a dual-luciferase reporter assay. Initially, the pmirGLO-circACACA-wild type plasmid (circACACA-WT), pmirGLO-circACACA-mutated type plasmid (circACACA-MT), pmirGLO-CBFB-wild type plasmid (CBFB-WT), and pmirGLO-CBFB-mutated type plasmid (CBFB-MT) were constructed Tsingke company (Beijing, China). These plasmids were then co-transfected with miRNA mimics or mimics NC into DF-1 cells. The Dual-Glo Luciferase Assay System Kits (Beyotime) were used to measure the activities of both Renilla and Firefly luciferases.

### Statistical analysis

All statistical analyses were conducted using SPSS 19.0 Statistics Software (SPSS, Chicago, IL, USA). The data were presented as the mean ± standard error of the mean (SEM). For comparisons between two groups, an unpaired Student’s t-test was utilized, while for multiple group comparisons, a one-way analysis of variance (ANOVA) was employed. Significant differences were denoted as follows: \**P* < 0.05, \*\**P* < 0.01. Additionally, different letters (a, b, c) were used to indicate statistical significance at *P* < 0.05.

## Results

### Overview of circRNAs deep sequencing data

High-throughput sequencing of circRNAs was conducted in both FL and CL samples. The raw data from six samples have been deposited in the SRA database with the accession number PRJNA960781. Between the two sample sets, circRNA-seq generated a total of 53,426,992 to 65,546,118 raw reads. After filtering out low-quality reads, the proportion of clean reads exceeded 98.18%, as detailed in S5 Table. The read density of each sample aligned to the chromosomes in the genome was statistically analyzed using circos, revealing that circRNAs are distributed across all chromosomes (S1A Fig).

CircRNAs were detected through a combined analysis involving find_circ and CIRI. In each sample, circRNAs were identified within exons, introns, and intergenic regions. Exonic circRNAs were the most abundant (Fig 1A; S6 Table), with lengths primarily ranging from 300 to 400 bp (S1B Fig). Across the six samples, a total of 3979 circRNAs were discovered (S7 Table). Among them, 3187 circRNAs were identified in FL, 2924 in CL, and 2132 were commonly expressed in both FL and CL (Fig 1B).

**Figure. 1.**
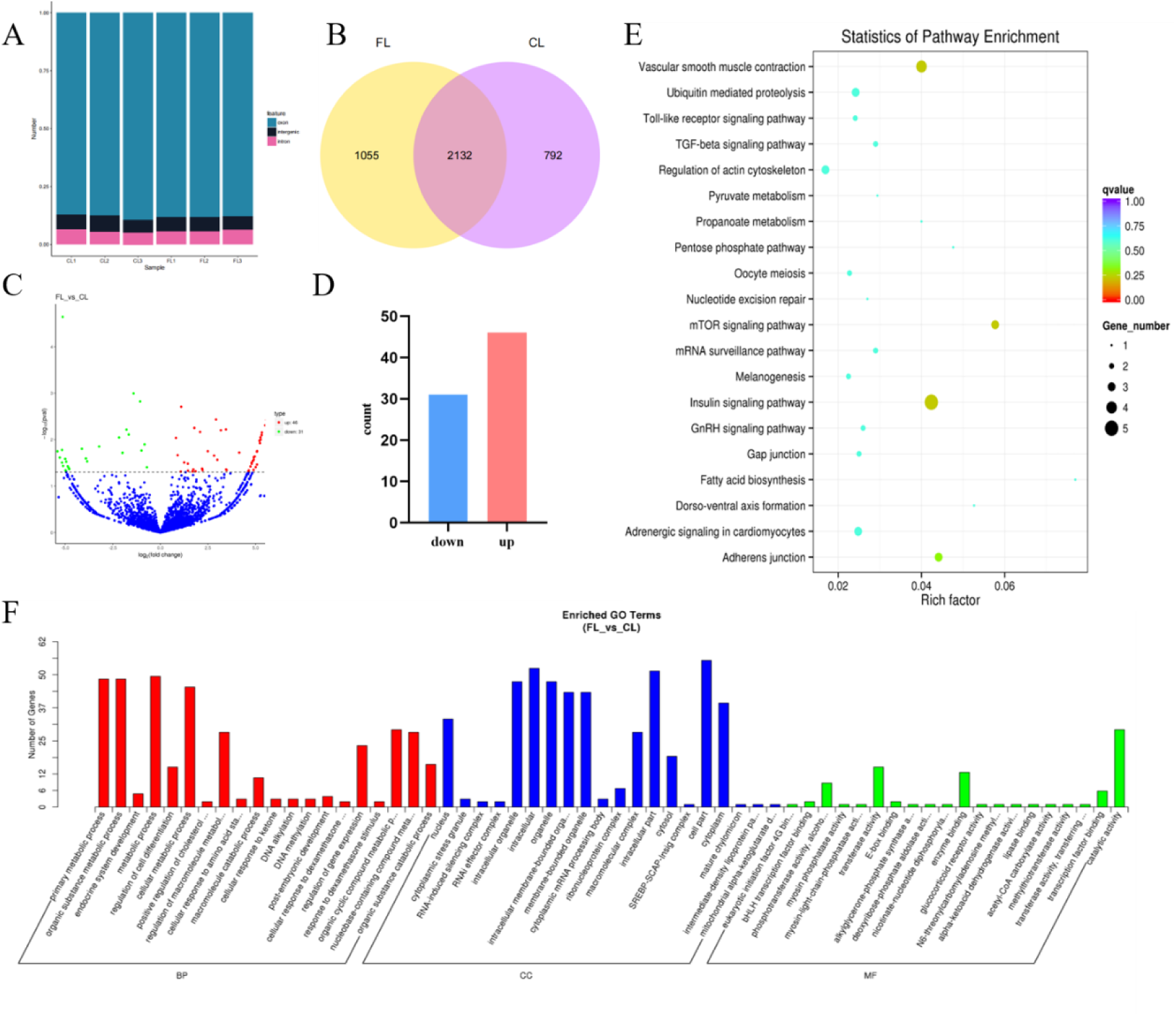
Overview of circRNA deep sequencing data and analysis of differentially expressed circRNAs. (A) Number of exons, introns and intergenic regions circRNAs per sample. (B) Validation of circRNAs within FL and CL. (C) Volcanic map of differentially expressed circRNAs in the CL and FL. (D) Analysis of differentially expressed circRNAs in the CL and FL. (E) KEGG enrichment for parental genes of differentially expressed circRNAs. (F) GO enrichment for the parental genes of differentially expressed circRNAs.

### The analysis of differentially expressed circRNAs

A total of 77 differentially expressed circRNAs were identified between FL and CL, with detailed information listed in S8 Table. Compared to CL, 46 circRNAs were upregulated in FL, while 31 were downregulated (Fig 1C and 1D). Cluster analysis revealed notable differences in circRNA expression patterns between the CL and FL groups of chickens (S1C Fig). Subsequently, GO and KEGG enrichment analysis were performed on the parental genes of the differentially expressed circRNAs. The KEGG analysis revealed that the genes were predominantly enriched in the insulin signaling pathway, mTOR signaling pathway, TGF-β signaling pathway, and metabolic pathways (Fig 1E). The GO results showed that the parental genes of the circRNAs were mainly associated with metabolic processes, cell parts, catalytic activity, and other categories (Fig 1F). Then, the circRNA-miRNA interaction network for the top 8 differentially highly expressed circRNAs were constructed and selected them as candidate genes for subsequent validation of biological functions (S9 Table; S1D Fig).

### Validation of differentially expressed circRNAs and selection of circACACA

The circular structure and expression patterns of these 8 differentially expressed circRNAs were verified. Initially, we analyzed the sequences of the circRNAs and designed divergent primers that spanned their splice sites. PCR amplification was then conducted, and the reaction products were subjected to Sanger sequencing to confirm the splice junctions (Fig 2A). Our results also indicated that random primers were more effective for amplifying circRNAs during reverse transcription compared to Oligo d(T) primers. Conversely, the opposite trend was observed for linear RNA (S1E Fig). To assess the stability of the circRNAs, RNase R treatment was used, and the results demonstrated that circRNAs exhibited greater resistance to RNase R compared to *β- actin* mRNA (Fig 2B). Finally, we validated the expression levels of these 8 circRNAs in FL and CL tissues using qPCR. The results revealed that circACACA, circKMT2E, circAHCTF1, circMAP4K5, circUSP45, and circRHEB were highly expressed in FL, whereas circPLXNC1 and circQPRT exhibited the opposite expression pattern (Fig 2C). Given its elevated expression level and notable differential expression between the FL and CL groups, we selected circACACA as the primary target circRNA for our experiment. Furthermore, we examined the expression pattern of circACACA across multiple chicken tissues. Our finding demonstrated that circACACA expression was markedly upregulated in the liver compared to other tissues (Fig 2D). To gain deeper insights into the intracellular localization of circACACA, we performed RNA-FISH analysis in both chicken hepatocytes and liver tissues. The results revealed that circACACA primarily resides in the cytoplasm and exhibits notably higher expression in the FL group (Fig 2E). These observations hint at a potentially pivotal regulatory role of circACACA in liver metabolism and development.

**Figure. 2.**
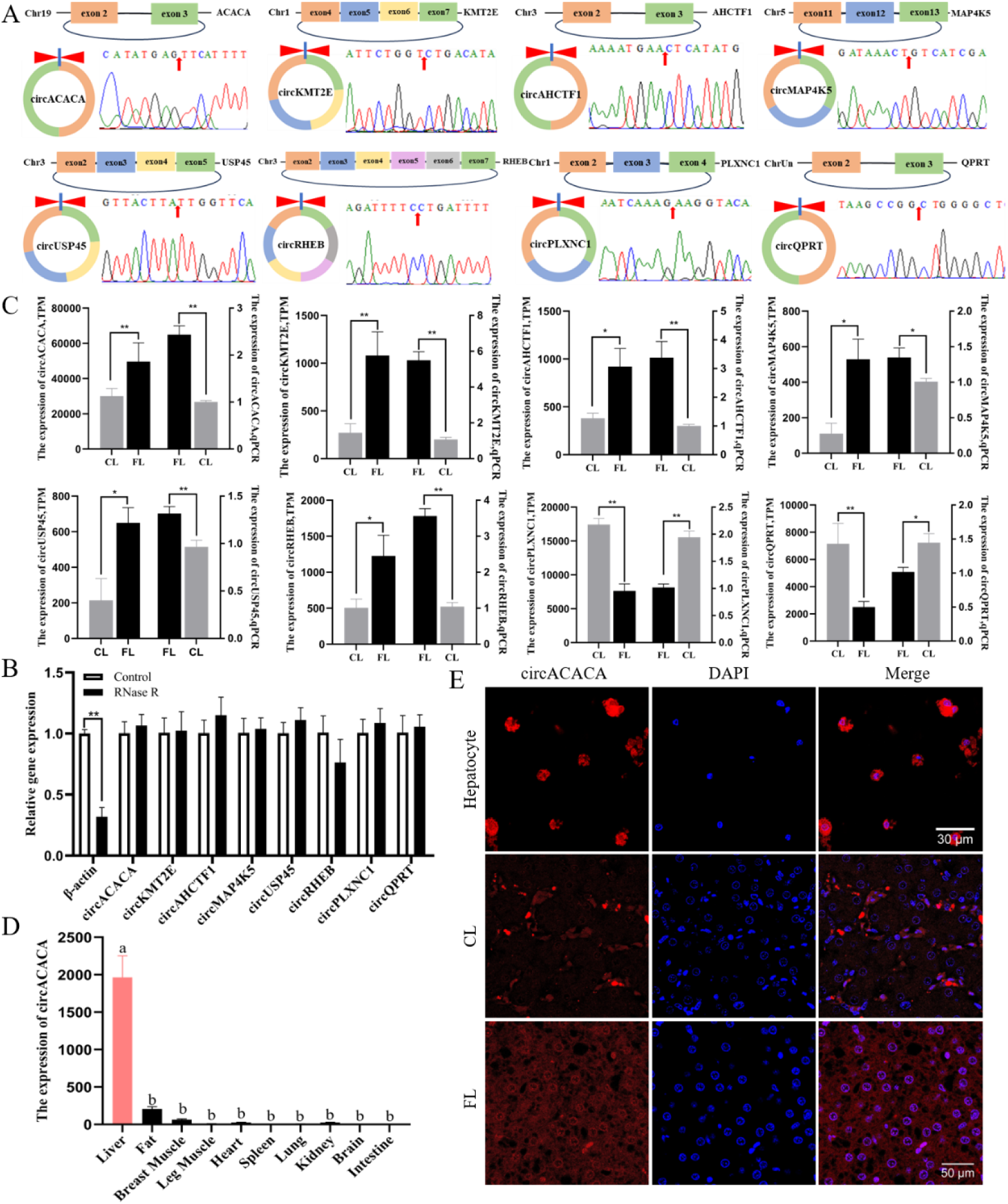
Validation of differentially expressed circRNAs and selection of circACACA. (A) Origin information of the top 8 differentially high-expressing circRNAs and Sanger sequencing validates the back-splicing junctions. Red arrows indicate splice sites. (B) RNase R analysis of digestion resistance for the top 8 differentially high-expressing circRNAs and *β-actin* in chicken hepatocytes. (C) Comparison of sequencing analysis and qPCR results of the top 8 differentially high-expressing circRNAs. (D) qPCR analysis of the mRNA level of circACACA in different tissues of chicken. (E) FISH analysis of the subcellular localization of circACACA in hepatocytes and liver tissue (FL and CL). Scale bars= 30 μm and 50 μm. The data is represented as mean ± SEM. * *P* < 0.05; ** *P* < 0.01.

### CircACACA damages liver healthy by promoting lipid accumulation, oxidative stress and autophagy in chicken hepatocytes

To explore the effects of circACACA on chicken liver metabolism, we successfully modulated its expression in hepatocytes by employing siRNA (si- circACACA) for knockdown and an overexpression vector (ov-circACACA) for upregulation (S1F and S1G Fig). The knockdown of circACACA resulted in a substantial downregulation of genes implicated in fat synthesis, namely *ACACA*, *SCD1*, *FASN*, and *SREBP* (S2A Fig). Conversely, it led to an upregulation of mRNA levels associated with lipid transport genes (*ApoVLDL-Ⅱ*, *APOA1*, and *CPT1*) (S2B Fig) and β-oxidation-related genes (*ACSL1*, *ACADS*, and *ACOX1*) (S2C Fig). Overexpression of circACACA exhibited converse effects on these gene expressions (S2D-S2F Fig). Similarly, protein expression analysis revealed a decrease in FASN and an increase in ACOX1 in the si-circACACA-treated group (Fig 3A; S2G Fig). In contrast, overexpression of circACACA significantly elevated FASN protein levels while decreasing ACOX1 (Fig 3B; S2H Fig). Furthermore, there was a marked accumulation of TG and TC in the ov-circACACA transfection group (Fig 3C and 3D). BODIPY 493/503 staining demonstrated a significant reduction in lipid droplet accumulation in hepatocytes upon circACACA knockdown (S2I and S2J Fig), whereas an extensive area of fluorescence staining was observed in the circACACA overexpression group (S2I and S2K Fig). These findings indicate that circACACA disrupts the balance of fat metabolism by promoting lipid accumulation in chicken livers.

**Figure. 3.**
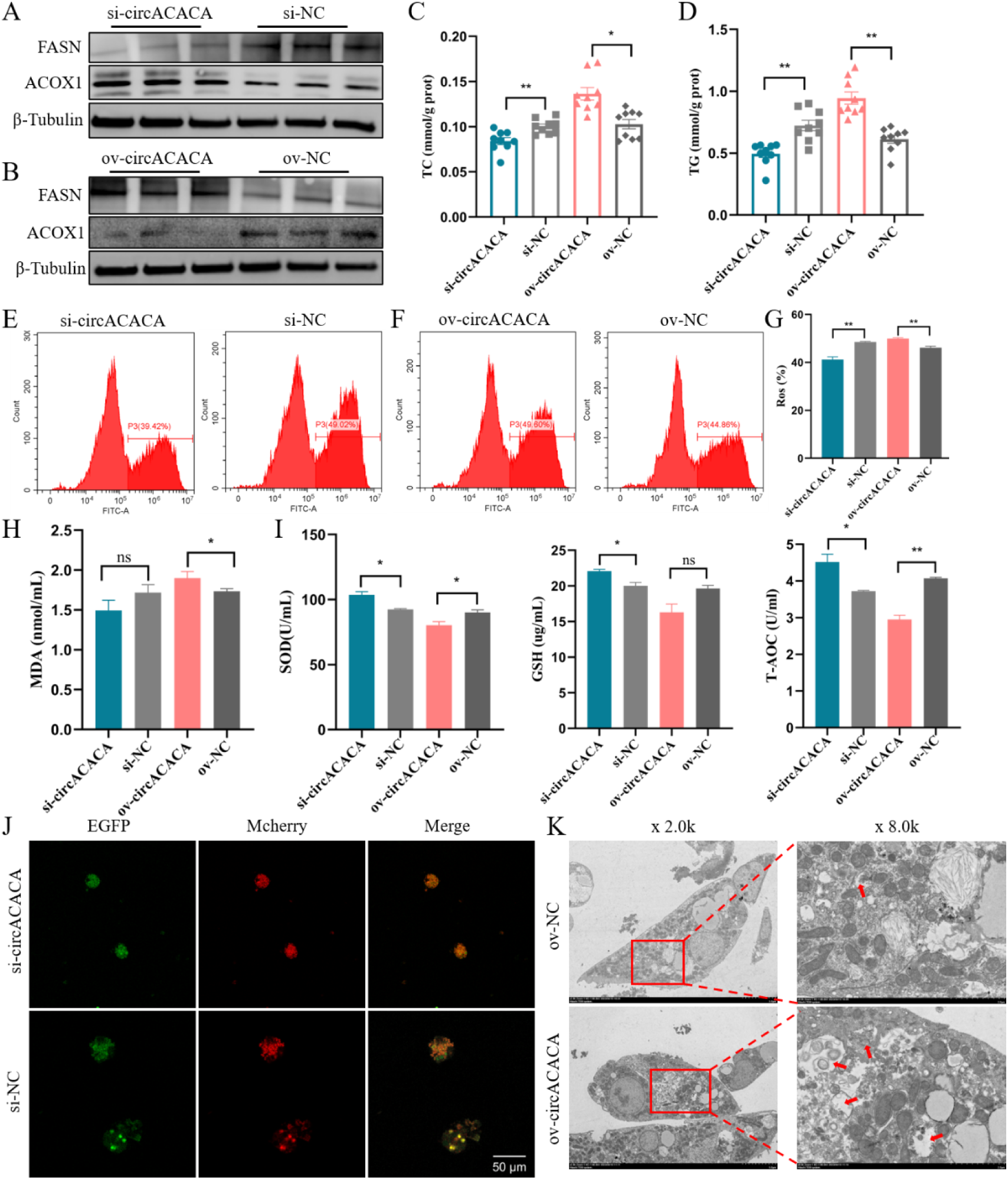
circACACA damages liver healthy by promoting lipid accumulation, oxidative stress and autophagy in chicken hepatocytes. (A, B) Western blot examination of the protein content of FSAN and ACOX1 after knockdown and overexpression of circACACA. (C, D) The content of TC and TG after silencing and increasing of circACACA. (E, F) Flow cytometry measurement of ROS levels after knockdown and upregulation of circACACA. (G) Measurement of intracellular ROS levels following circACACA expression modulation (H) The level of intracellular MDA after interference and increasing of circACACA. (I) The content of intracellular SOD, GSH and T-AOC after changing the expression of circACACA. (J) Adenovirus fluorescence image of Mcherry-EGFP-LC3 after interference of circACACA. Mcherry was employed to label and track LC3. Scale bars= 50 μm. (K) Transmission electron micrograph of hepatocytes after overexpression of circACACA. The red arrow shows the position of autophagolysosomes. The data is represented as mean ± SEM. * *P* < 0.05; ** *P* < 0.01.

To elucidate the underlying role of circACACA on antioxidant capacity, we conducted comprehensive analyses of the mRNA and protein expression of genes implicated in antioxidant stress responses. qPCR analysis demonstrated a significant upregulation of *SOD*, *GPX*, *TRX*, and *GST* mRNA levels upon circACACA knockdown (S3A Fig). Conversely, overexpression of circACACA resulted in a reduction in the mRNA expression of these genes (S3B Fig). Consistent with the mRNA changes, alterations in circACACA expression were mirrored by corresponding changes in the protein levels of SOD and GPX7 (S3C and S3D Fig). Flow cytometric analysis of intracellular ROS revealed a substantial decrease in ROS production following circACACA interference compared to the control group (Fig 3E and 3G). Conversely, overexpression of circACACA had the converse effect, leading to an increase in ROS levels (Fig 3F and 3G). Furthermore, there was a positive correlation between circACACA expression and the accumulation of MDA (Fig 3H), while it exhibited an inverse correlation with the production of antioxidant enzymes such as SOD, GSH, and T-AOC (Fig 3I). Collectively, these findings establish that circACACA impairs the antioxidant stress capacity in chicken hepatocytes.

Subsequently, we explored the biological function of circACACA in modulating autophagy in chicken hepatocytes. Our findings revealed a significant downregulation of mRNA expression for *Beclin1*, *ATG5*, *ATG7*, and *ATG9* following circACACA interference, concomitant with an upregulation of *P62* expression (S3E Fig). Silencing of circACACA led to a reduction in the levels of LC3-II and Beclin1 proteins, while elevating P62 levels (S3F Fig). Conversely, the alterations in mRNA and protein expression levels of autophagy-related genes observed upon overexpression of circACACA were diametrically opposed to those seen after circACACA knockdown (S3G and S3H Fig). Furthermore, we utilized Mcherry-EGFP-LC3 adenovirus as a tool to monitor the autophagy marker LC3. As illustrated in Fig 3J, a marked decrease in the number of autophagosomes (Merge) was evident upon circACACA knockdown. Electron microscopy analysis provided additional support, demonstrating that circACACA stimulates the formation of autophagosomes in hepatocytes (Fig 3K). Taken together, these results compellingly suggest that circACACA enhances autophagy in chicken hepatocytes.

### CircACACA acts as a ceRNA for miR-132b-5p and miR-101-2-5p

Utilizing RNAhybrid software, we predicted potential target miRNAs for circACACA and conducted a Venn analysis in conjunction with circRNA sequencing data. Our analysis identified 9 commonly predicted miRNAs: miR-132b-5p, miR-1653, miR-301b-5p, miR-6596-3p, miR-130b-5p, miR-3531-3p, miR-1771, miR-1626-5p, and miR-101-2-5p (S3I Fig). Subsequent preliminary validation using qPCR revealed that only miR-132b-5p and miR-101-2-5p exhibited expression levels inversely correlated with circACACA (Fig 4A and 4B; S3J Fig). As depicted in Fig 4C and 4D, the potential binding sites between circACACA and these two miRNAs are illustrated. To further validate their target relationship, we conducted RNA-FISH experiments to assess their subcellular localization within hepatocytes. Our results demonstrated that circACACA co-localized with both miR-132b-5p and miR-101-2-5p in hepatocytes (Fig 4E and 4F). Additionally, we employed a dual-luciferase reporter assay utilizing circACACA-WT and circACACA-MT plasmids, which contained either normal or mutated binding sites for miR-132b-5p and miR-101-2-5p in the 5’ seed region (Fig 4G and 4H). The results of this assay revealed that both miR-132b-5p and miR-101-2-5p significantly reduced the luciferase activity of the circACACA-WT plasmid compared with the circACACA-MT plasmid (Fig 4I and 4J). Collectively, these findings provide compelling evidence that circACACA functions as a molecular sponge for miR-132b- 5p and miR-101-2-5p.

**Figure. 4.**
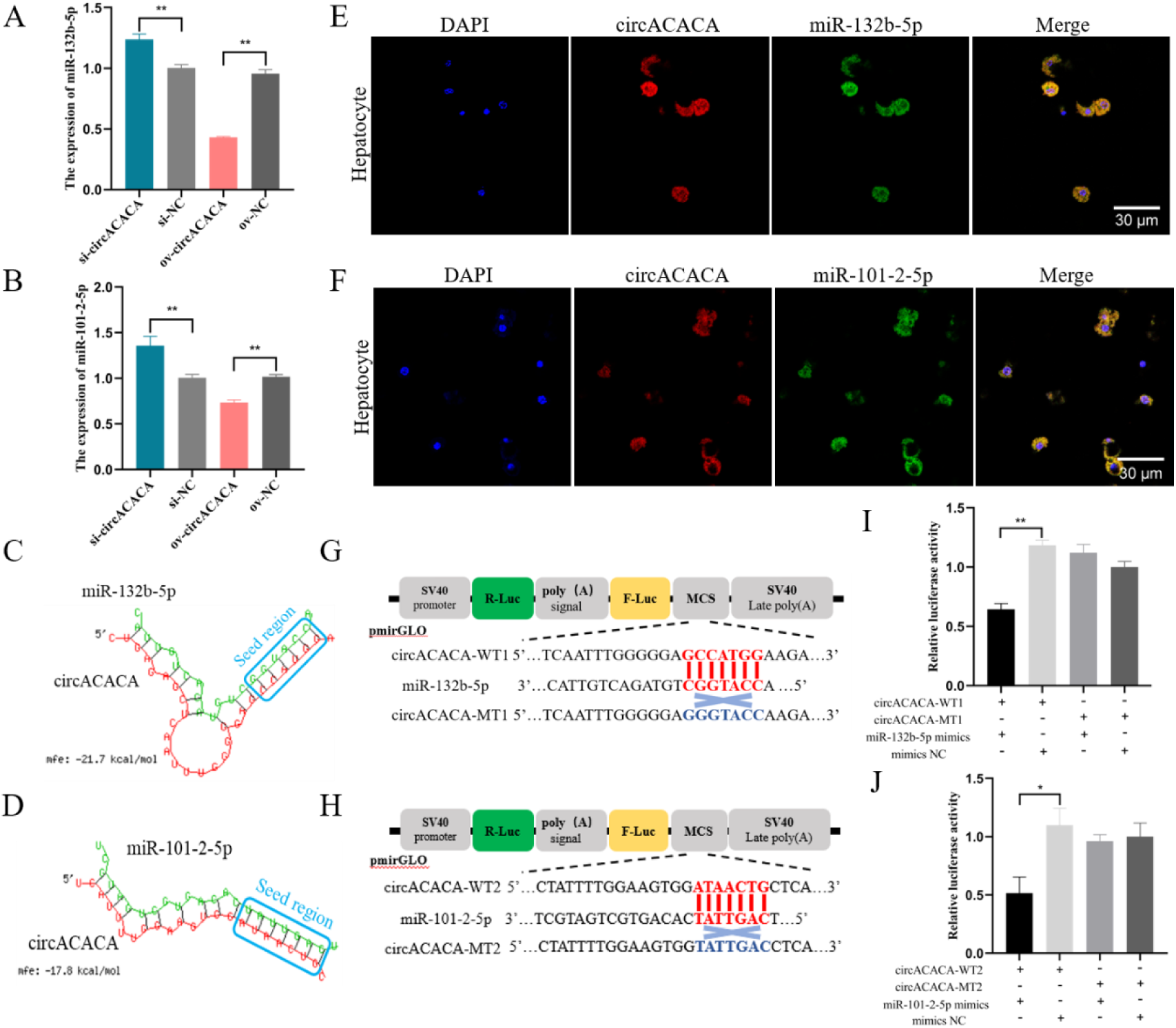
circACACA acts as a ceRNA for miR-132b-5p and miR-101-2-5p. (A, B) qPCR examination of the expression of miR-132b-5p and miR-101-2-5p after interference and overexpression of circACACA. (C, D) The RNAhybrid software predicted binding sites between circACACA and miR-132b-5p/miR-101-2-5p. (E, F) FISH examination of the targeting relationship between circACACA and miR-132b-5p/miR-101-2-5p in chicken hepatocytes. Scale bars= 30 μm. (G, H) Construction of wild type dual-luciferase reporter vector (circACACA-WT1, circACACA-WT2) and mutant type dual-luciferase reporter vector (circACACA-MT1, circACACA-MT2) of circACACA containing miR-132b-5p or miR-101- 2-5p binding sites. The seed sequences were highlighted in red, while the mutant sequences were marked in blue. (I, J) The dual-luciferase reporter assay of the relative luciferase activity after co-transfection with circACACA-WT or circACACA-MT and mimics miR-132b- 5p/miR-101-2-5p or mimics NC. The data is represented as mean ± SEM. * *P* < 0.05; ** *P* < 0.01.

### MiR-132b-5p and miR-101-2-5p contribute to the health of the liver by mitigating fat accumulation, oxidative damage and autophagy in chicken hepatocytes

Firstly, we investigated the potential impact of miR-132b-5p and miR-101-2-5p on liver fat metabolism. qPCR results revealed that the mRNA levels of fat synthesis genes were upregulated upon the reduction of miR-132b-5p, while lipid transport-related genes and β-oxidation-related genes were suppressed (S4A Fig). Conversely, the elevation of exogenous miR-132b-5p had the opposite effect on the expression of these genes (S4B Fig). Western blot analysis further demonstrated that miR-132b-5p increased the protein expression of ACOX1 while decreasing the protein level of FASN (Fig 5A and 5B; S4C and S4D Fig). Additionally, miR-132b-5p reduced the accumulation of intracellular TG and total TC (Fig 5C). Through BODIPY 493/503 staining experiments, we observed a large area of fluorescence staining in the miR-132b-5p inhibitor group (S4E and S4F Fig). However, compared to the miR-132b-5p mimics-treated group, the accumulation of lipid droplets in the cells significantly decreased upon the increase in miR-132b-5p expression (S4E and S4G Fig). Similarly, miR-101-2-5p exhibited comparable effects on the mRNA expression of these genes as miR-132b-5p (S5A and S5B Fig). Likewise, miR-101-2-5p elevated the protein level of ACOX1 and reduced FASN (Fig 5E and 5F; S5C and S5D Fig), and the downregulation of miR-101-2-5p led to an increase in TG and TC content (Fig 5D). The inhibition of miR-101-2-5p notably induced lipid content in hepatocytes (S5E and S5F Fig), while the elevation of miR-101-2-5p significantly reduced the staining area (S5E and S5G Fig). Collectively, these results suggest that both miR-132b-5p and miR- 101-2-5p reduce the accumulation of lipid droplets in chicken hepatocytes.

**Figure. 5.**
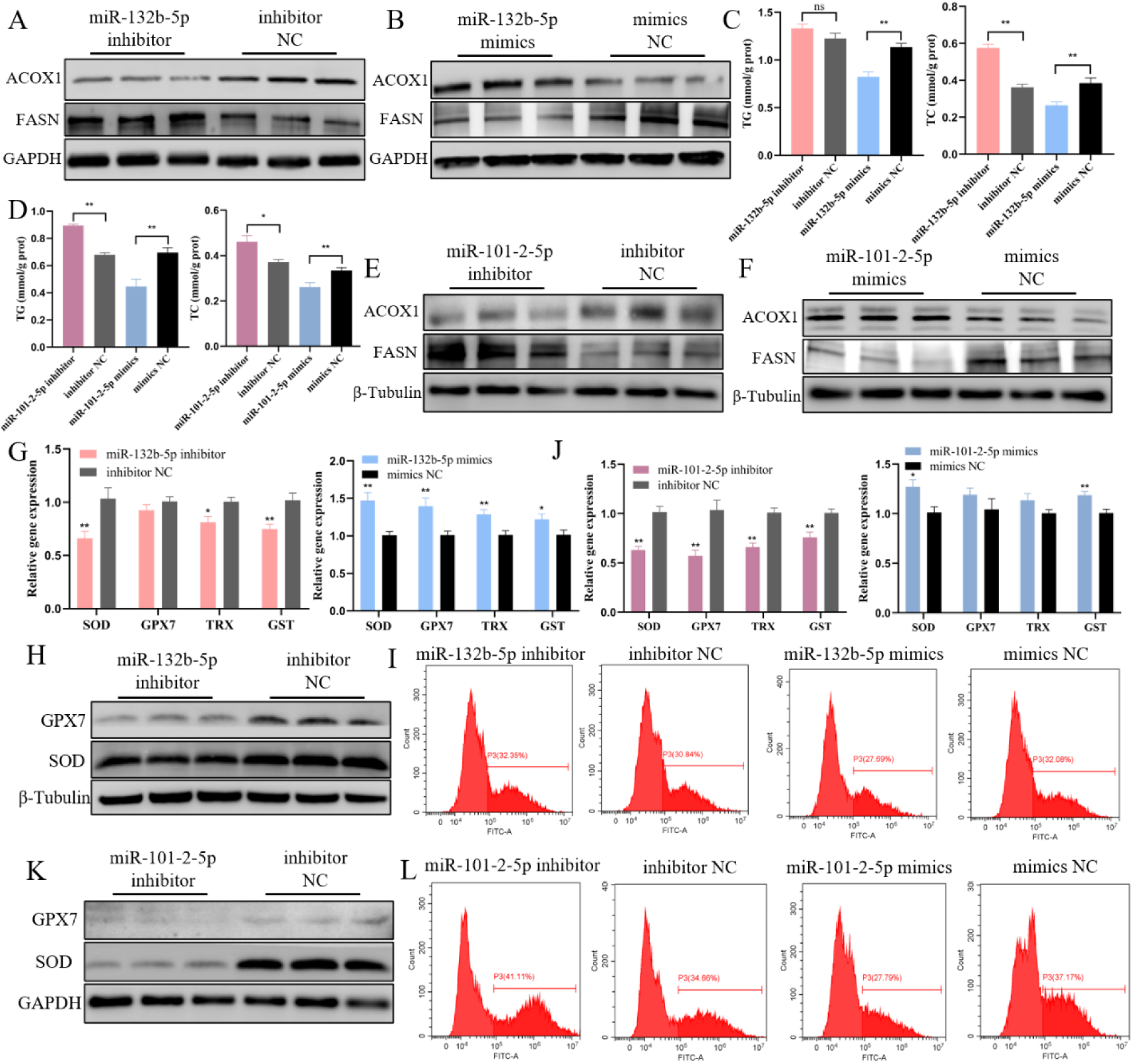
miR-132b-5p and miR-101-2-5p contribute to the health of the liver by mitigating fat accumulation, oxidative damage and autophagy. (A, B) Western blot assessment of the protein expression of ACOX1 and FASN after interference and overexpression of miR-132b-5p. (C) The content of TG and TC after changing of miR-132b- 5p in chicken hepatocytes. (D) The content of TG and TC after changing of miR-101-2-5p in chicken hepatocytes. (E, F) Western blot measurement of the protein expression of ACOX1 and FASN after knockdown and overexpression of miR-101-2-5p. (G) qPCR analysis of the mRNA level of anti-oxidative stress related genes after changing miR-132b-5p. (H)Western blot assessment of the protein expression of GPX7 and SOD after knockdown of miR-132b-5p. (I) Flow cytometry analysis of ROS levels after knockdown and overexpression of miR-132b- 5p. (J) qPCR analysis of the mRNA level of anti-oxidative stress related genes after the decreasing and increasing of miR-101-2-5p. (J) Western blot assessment of the protein expression of GPX7 and SOD after downregulation of miR-101-2-5p. (L) Flow cytometry analysis of ROS levels after knockdown and overexpression of miR-101-2-5p. The data is represented as mean ± SEM. * *P* < 0.05; ** *P* < 0.01.

Subsequently, we discovered that inhibiting miR-132b-5p significantly decreased the mRNA levels of *SOD*, *TRX*, and *GST*, whereas their expression was augmented upon miR-132b-5p overexpression (Fig 5G). Furthermore, the protein levels of SOD and GPX7 were downregulated in the miR-132b-5p inhibitor-transfected group (Fig 5H; S6A Fig), while overexpression of miR-132b-5p promoted their protein expression (S6B Fig). Additionally, ROS activity increased following the downregulation of miR- 132b-5p, while its overexpression had the opposite effect (Fig 5I; S6C Fig). Moreover, we found that miR-132b-5p reduced the levels of MDA (S6D Fig) while increasing the concentrations of SOD, GSH, and T-AOC (S6E-S6G Fig). Similarly, miR-101-2-5p also elevated the expression of genes related to antioxidant stress (Fig 5J). Upon altering the expression of miR-101-2-5p, the changes in the protein concentrations of SOD and GPX7 mirrored their mRNA level alterations (Fig 5K; S6H and 6I Fig). Likewise, miR-101-2-5p also attenuated the accumulation of ROS within the cells (Fig 5L; S6J Fig). MiR-101-2-5p exhibited comparable effects on MDA accumulation (S6K Fig) and antioxidant enzyme activity (S6L-S6N Fig). Collectively, these results demonstrate that both miR-132b-5p and miR-101-2-5p alleviate oxidative stress in chicken hepatocytes.

Downregulation of miR-132b-5p resulted in elevated mRNA and protein expression of autophagy-related genes (Fig 6A and 6B; S6O Fig), which were subsequently suppressed upon overexpression of miR-132b-5p (excluding P62) (Fig 6C; S6P and S6Q Fig). The modulation of miR-101-2-5p led to an increase in the mRNA levels of autophagy-related genes (Fig 6D) and the protein levels of LC3-II and Beclin1 (but not P62) (Fig 6E; S6R Fig). Conversely, overexpression of miR-101-2-5p negatively correlated with the mRNA and protein levels of these autophagy-related genes (Fig 6F; S6S and S6T Fig). The Mcherry-EGFP-LC3 staining assay revealed a significant decrease in the area of autophagosomes in the miR-132b-5p mimics treatment group (Fig 6G), and electron microscopy further confirmed the presence of a large number of autolysosomes in the miR-132b-5p inhibitor group (Fig 6H). Similarly, the formation of autophagosomes was markedly reduced upon overexpression of miR- 101-2-5p (Fig 6I), whereas it was enhanced in the miR-101-2-5p inhibitor group (Fig 6J). Taken together, these results confirm that both miR-132b-5p and miR-101-2-5p possess the ability to attenuate autophagy in chicken hepatocytes.

**Figure. 6.**
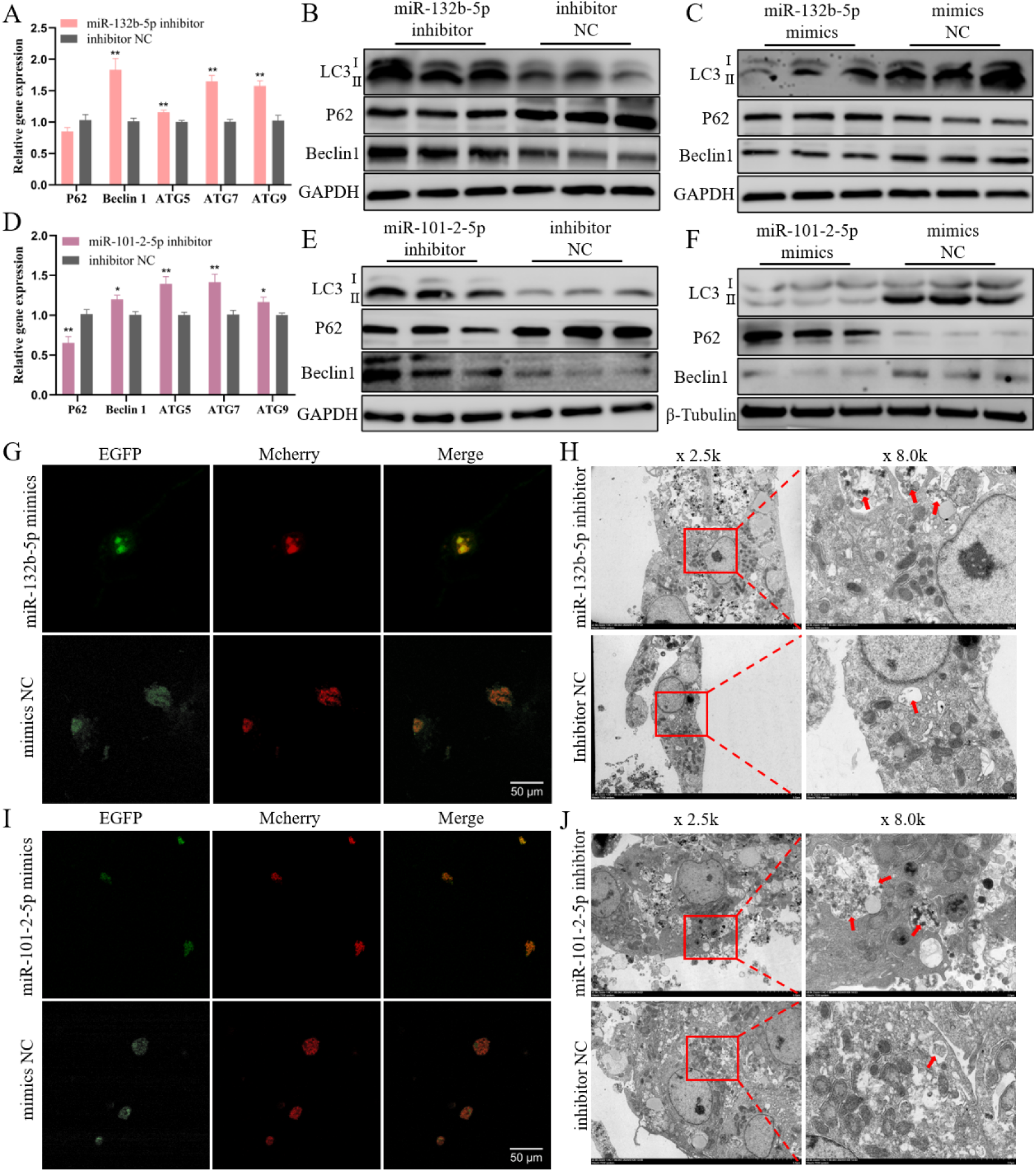
miR-132b-5p and miR-101-2-5p reduces autophagy of chicken hepatocytes. (A) qPCR measurement of the mRNA level of autophagy related genes after inhibition of miR- 132b-5p. (B, C) Western blot assessment of the protein content of autophagy related genes after alter the expression of miR-132b-5p. (D) qPCR analysis of the mRNA expression of autophagy related genes after interference of miR-101-2-5p. (E, F) Western blot evaluation of the protein concentration of autophagy related genes after altering the expression of miR-101-2-5p. (G) Fluorescence image of Mcherry-EGFP-LC3-labeled chicken hepatocytes after overexpression of miR-132b-5p. Scale bars= 50 μm. (H) Transmission electron micrograph of hepatocytes after inhibition of miR-132b-5p. The red arrow highlights auto-phagolysosomes. (I) Image of Mcherry-EGFP-LC3 fluorescence after the upregulation of miR-101-2-5p. (J) Transmission electron micrograph of hepatocytes following knockdown of miR-101-2-5p. The red arrow highlights auto-phagolysosomes. The data is represented as mean ± SEM. * *P* < 0.05; ** *P* < 0.01.

### CBFB is a common target of miR-132b-5p and miR-101-2-5p

To investigate the regulatory mechanisms associated with miR-132b-5p and miR- 101-2-5p, we employed two online prediction tools, miRDB and TargetScan, alongside RNA-seq data to identify potential target genes for these two miRNAs (S10 Table). Venn analysis subsequently revealed that CBFB is a common target gene for both miR- 132b-5p and miR-101-2-5p (Fig 7A). The specific binding sites of these two miRNAs to CBFB were also predicted (Fig 7B). The qPCR experiments demonstrated that increased expression of miR-132b-5p and miR-101-2-5p led to decreased CBFB expression, whereas reduced expression of these two miRNAs correlated with increased CBFB expression (Fig 7C and 7D). Next, we constructed CBFB-WT1 and CBFB-MT1 vectors (Fig 7E), as well as CBFB-WT2 and CBFB-MT2 vectors (Fig 7F). Using dual-luciferase reporter assays, we assessed changes in fluorescent enzyme activity. The results revealed that the addition of miR-132b-5p mimics and miR-101-2- 5p mimics significantly reduced the fluorescent activity of CBFB-WT1 and CBFB- WT2 (Fig 7G and 7H). Collectively, these results suggest that CBFB is a common target of both miR-132b-5p and miR-101-2-5p.

**Figure. 7.**
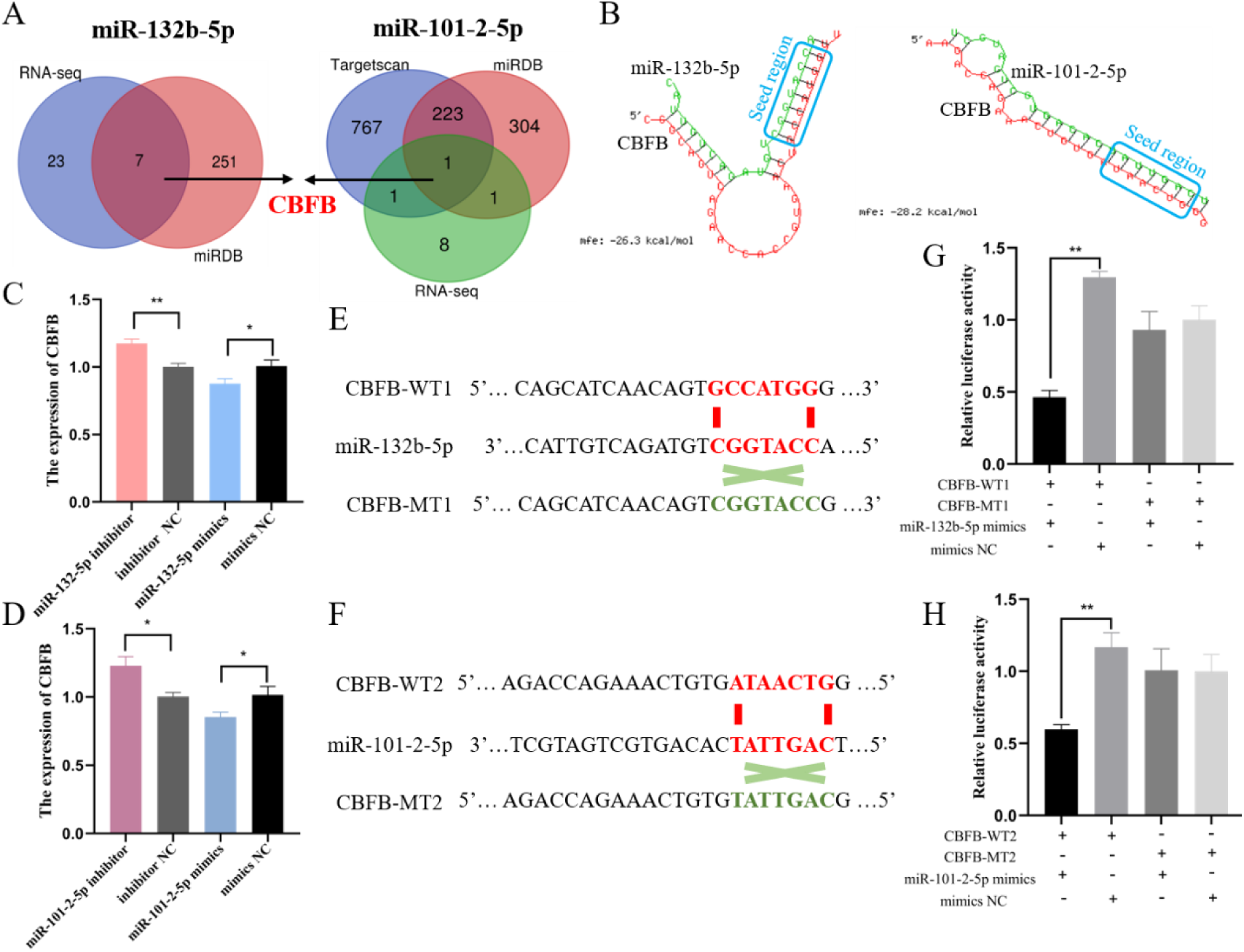
CBFB is a common target of miR-132b-5p and miR-101-2-5p. (A) The Venn diagram of miR-132b-5p and miR-101-2-5p target prediction genes. (B) The predicted interaction sites between CBFB and miR-132b-5p or miR-101-2-5p. (C, D) qPCR analysis of the mRNA level of CBFB after interference and overexpression of miR-132b-5p and miR-101- 2-5p. (E, F) Construction of wild type dual-luciferase reporter vector (CBFB-WT1, CBFB- WT2) and mutant type dual-luciferase reporter vector (CBFB-MT1, CBFB-MT2) of CBFB containing miR-132b-5p or miR-101-2-5p binding sites. The seed sequences were colored red, and the mutant sequences were colored green. (G, H) The dual-luciferase reporter analysis of the relative luciferase activity after co-transfection with CBFB-WT or CBFB-MT and mimics miR-132b-5p/miR-101-2-5p or mimics NC. The data is represented as mean ± SEM. * *P* < 0.05; ** *P* < 0.01.

### CBFB impairs liver health by promoting fat accumulation, oxidative damage and autophagy in chicken hepatocytes

We investigated the potential effects on chicken liver health by modulating CBFB expression in vitro (S7A and S7B Fig). The knockdown of CBFB resulted in a significant reduction in the mRNA levels of *ACACA*, *SCD1*, *FASN*, and *SREBP* (S7C Fig), while concurrently increasing the expression of genes related to lipid transport and β-oxidation (S7D and S7E Fig). Conversely, the overexpression of CBFB had the opposite effect on their expression (Fig S7F-S7H Fig). Western blot analysis yielded results consistent with those obtained from qPCR analysis (Fig S7I-S7L Fig). BODIPY 493/503 staining revealed that CBFB knockdown significantly decreased lipid droplet accumulation compared to the control group, whereas the ov-CBFB group exhibited extensive fluorescence staining (Fig 8A; S7M Fig). Furthermore, we observed a significant increase in TG and TC levels following the upregulation of CBFB (Fig 8B and 8C). Collectively, our findings suggest that CBFB promotes fat accumulation in chicken hepatocytes.

**Figure. 8.**
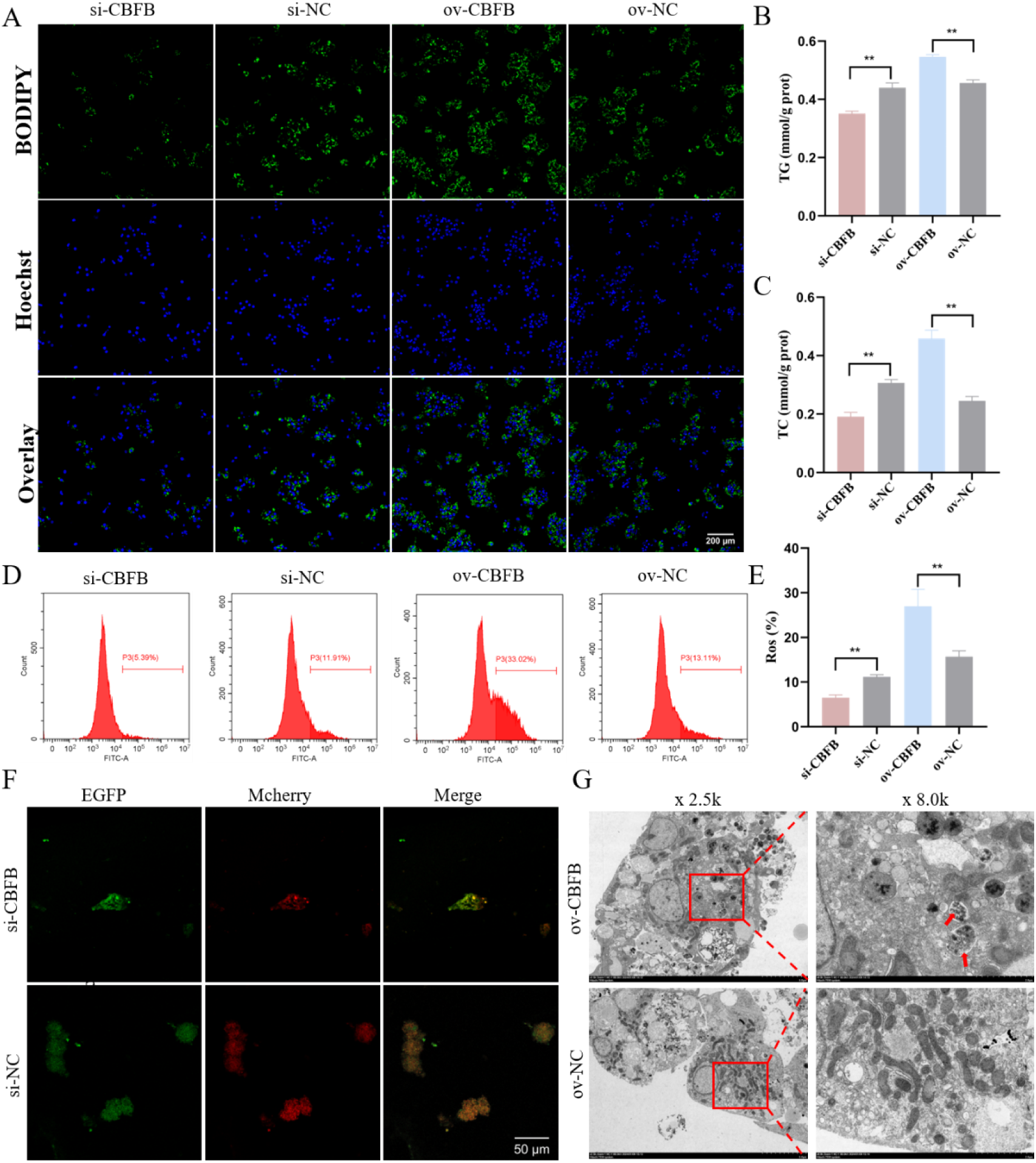
CBFB impairs liver health by promoting fat accumulation, oxidative damage and autophagy. (A) BODIPY 493/503 staining of lipid droplets after the interference and overexpression of CBFB. Scale bars= 200 μm. (B, C) The content of intracellular TG and TC after knockdown and overexpression of CBFB. (D, E) Flow cytometric assessment of ROS levels upon downregulation and upregulation of CBFB. (F) Fluorescence microscopy of Mcherry-EGFP-LC3 after blocking CBFB. Scale bars= 50 μm. (G) Transmission electron microscopy image of hepatocytes following CBFB overexpression. The data is represented as mean ± SEM. * *P* < 0.05; ** *P* < 0.01.

The modulation of CBFB expression exerted a profound impact on the antioxidant defense system in chicken hepatocytes. Specifically, the knockdown of CBFB led to a notable upregulation of mRNA levels for *SOD*, *GPX7*, and *GST* (Fig S7N Fig). Conversely, elevated CBFB expression resulted in the suppression of these antioxidant genes (Fig S7O Fig). Consistent with the mRNA data, alterations in CBFB expression were mirrored at the protein levels for both GPX7 and SOD (Fig S7P and S7Q Fig). Furthermore, CBFB significantly augmented the production of ROS within the hepatocytes (Fig 8D and 8E). Collectively, these findings indicate that CBFB, analogous to circACACA, promotes oxidative stress in chicken hepatocytes.

After modulating CBFB expression in vitro, we observed corresponding changes in the mRNA levels of autophagy-related genes (S7R and S7S Fig). In the si-CBFB treatment group, the protein levels of LC3-II and Beclin1 were decreased, while the level of P62 was increased (S7T Fig). Conversely, the overexpression of CBFB had the opposite effect on the protein expression of these markers (S7U Fig). Mcherry-EGFP- LC3 staining revealed minimal fluorescence only in the CBFB-interfered cells (Fig 8F). Electron microscopy further confirmed these findings, showing a higher number of autophagosomes in the ov-CBFB group compared to the ov-NC group (Fig 8G). In summary, our results demonstrate that CBFB promotes autophagy in chicken hepatocytes.

### CircACACA inactivates AKT/mTOR and PPAR-γ signaling pathways by targeting miR-132b-5p and miR-101-2-5p to regulate CBFB/PIM1 complexes

We further delved into the mechanism underlying CBFB’s action. Western blot analysis revealed that CBFB did not alter the total protein expression of AKT and mTOR; however, upon CBFB knockdown, their phosphorylation levels were significantly elevated. Concurrently, the protein levels of markers of the PPAR-γ signaling pathway, namely PPAR-γ and RXRA, were also increased (Fig 9A; S8A Fig). Conversely, CBFB overexpression led to a notable decrease in the protein concentrations of p-AKT and p-mTOR, along with a reduction in PPAR-γ pathway activity (Fig 9B; S8B Fig). Additionally, we co-transfected hepatocytes with ov- circACACA, specific miRNAs, and si-CBFB. CBFB knockdown potentiated the inhibitory effect of circACACA on the phosphorylation levels of AKT and mTOR, as well as the protein expression of RXRA and PPAR-γ (Fig 9C; S8C Fig). CircACACA was found to interact with CBFB to regulate fat metabolism (as evidenced by changes in FASN and ACOX1), oxidative stress (SOD and GPX7), and autophagy (P62 and Beclin1) (Fig 9C; S8D Fig).

**Figure. 9.**
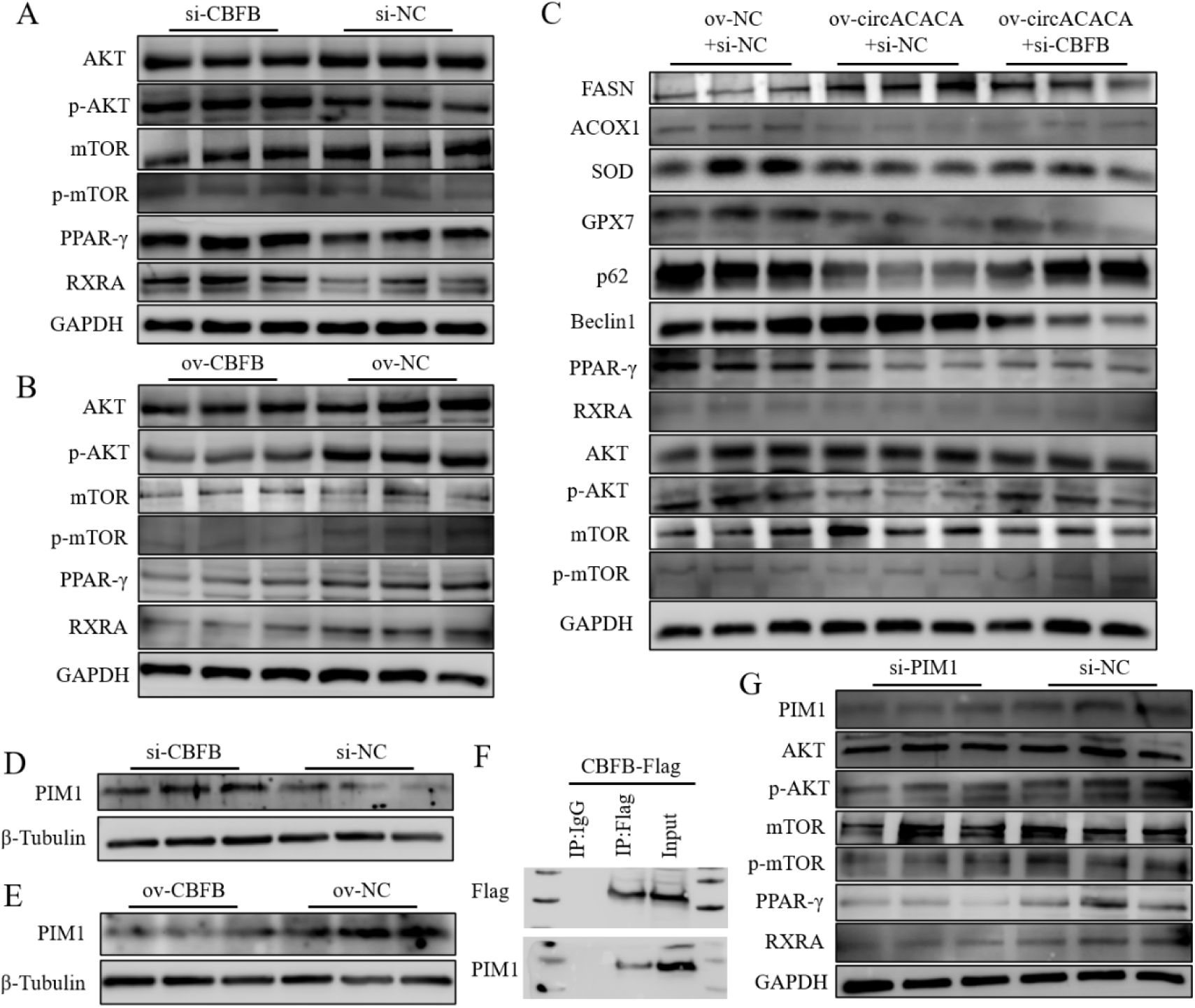
CircACACA inactivates AKT/mTOR and PPAR-γ signaling pathways by targeting miR-132b-5p and miR-101-2-5p to regulate CBFB/PIM1 complexes. (A, B) Western blot assessment of the levels of key protein in the AKT/mTOR (AKT, p-AKT, mTOR, p-mTOR) and the PPAR-γ (PPAR-γ and RXRA) signaling pathway after knockdown and overexpression of CBFB. (C) Western blot evaluation of the contents of fat metabolism related protein, anti-oxidative stress related protein, autophagy related protein, the AKT/mTOR and the PPAR-γ signaling pathway key protein after co-transfection with ov-NC+si-NC and ov- circACACA+si-NC or si-CBFB. (D, E) Western blot analysis of the protein level of PIM1 after altering CBFB expression. (F) Co-Immunoprecipitation (Co-IP) validation of the interaction between PIM1 and CBFB. (G) Western blot evaluation of the contents of key protein in the AKT/mTOR and the PPAR-γ signaling pathway after interference of PIM1. The data is represented as mean ± SEM. * *P* < 0.05; ** *P* < 0.01.

Subsequently, we validated the binding interaction between CBFB and PIM1 in chicken liver. Initially, we observed that the mRNA expression levels of PIM1 and CBFB displayed opposing patterns (S8E Fig). Western blot analysis revealed a significant increase in PIM1 protein content upon CBFB downregulation, whereas PIM1 levels were reduced following CBFB overexpression (Fig 9D and 9E; S8F Fig). Co-immunoprecipitation (Co-IP) results further confirmed this interaction, as both Flag and PIM1 proteins were detected in the input and Flag groups, but not in the IgG group (Fig 9F). Additionally, western blot analysis demonstrated that PIM1 knockdown reduced the activity of the AKT/mTOR and PPAR-γ signaling pathways (Fig 9G; S8G Fig). To further explore this relationship, co-transfection experiments were conducted, and revealed that downregulation of PIM1 expression could reverse the effects of si- CBFB on the AKT/mTOR and PPAR-γ pathways (S8H-S8J Fig). Furthermore, our results indicated that the introduction of circACACA reversed the effects of miR-132b- 5p and miR-101-2-5p on functional proteins, as well as on the AKT/mTOR signaling pathway and PPAR-γ pathway (S9A-S9F Fig). These findings collectively suggest that CBFB and PIM1 synergistically regulate the AKT/mTOR and PPAR-γ signaling pathways in the chicken liver. In summary, our collective findings indicate that the circACACA-miR-132b-5p/miR-101-2-5p-CBFB regulatory axis disrupts liver metabolism and development by inhibiting the AKT/mTOR and PPAR-γ pathways through its interaction with PIM1.

### Conservation analysis of circACACA and validation of the circularity and expression pattern of mmu-circAcaca

Initially, we retrieved and analyzed the circAtlas database to investigate the sequence characteristics and conservation of circACACA in humans, mice, and chickens. Our analysis revealed that circACACA exhibits similar exon numbers and positions across these species, with a high sequence similarity of 72.67% to the human sequence and 69.67% to the mouse sequence (S11 Table). Subsequently, we selected C57BL/6J mice as our model animals to validate the homology of circRNAs. We established a mouse model of FL by feeding the mice a high-fat diet and confirmed the successful induction of the model through various criteria. Morphologically, the FL mice were visibly larger and exhibited obesity compared to the control group. Upon dissection, their livers appeared yellowish, swollen, and loose (Fig 10A). Oil Red O staining demonstrated a significant increase in the area of red-positive staining in the high-fat treatment group, and HE staining further confirmed the accumulation of lipid droplets within liver cells (Fig 10B). Additionally, serum levels of TG and TC were significantly higher in the FL group compared to the CL group (ided additional evidence supporting the successful establishment of the mouse model of FL (S10B Fig). To validate the functional homology of circACACA, we utilized the AML12 mouse liver cell line to assess the stability and conservation of mmu-circAcaca.

**Figure. 10.**
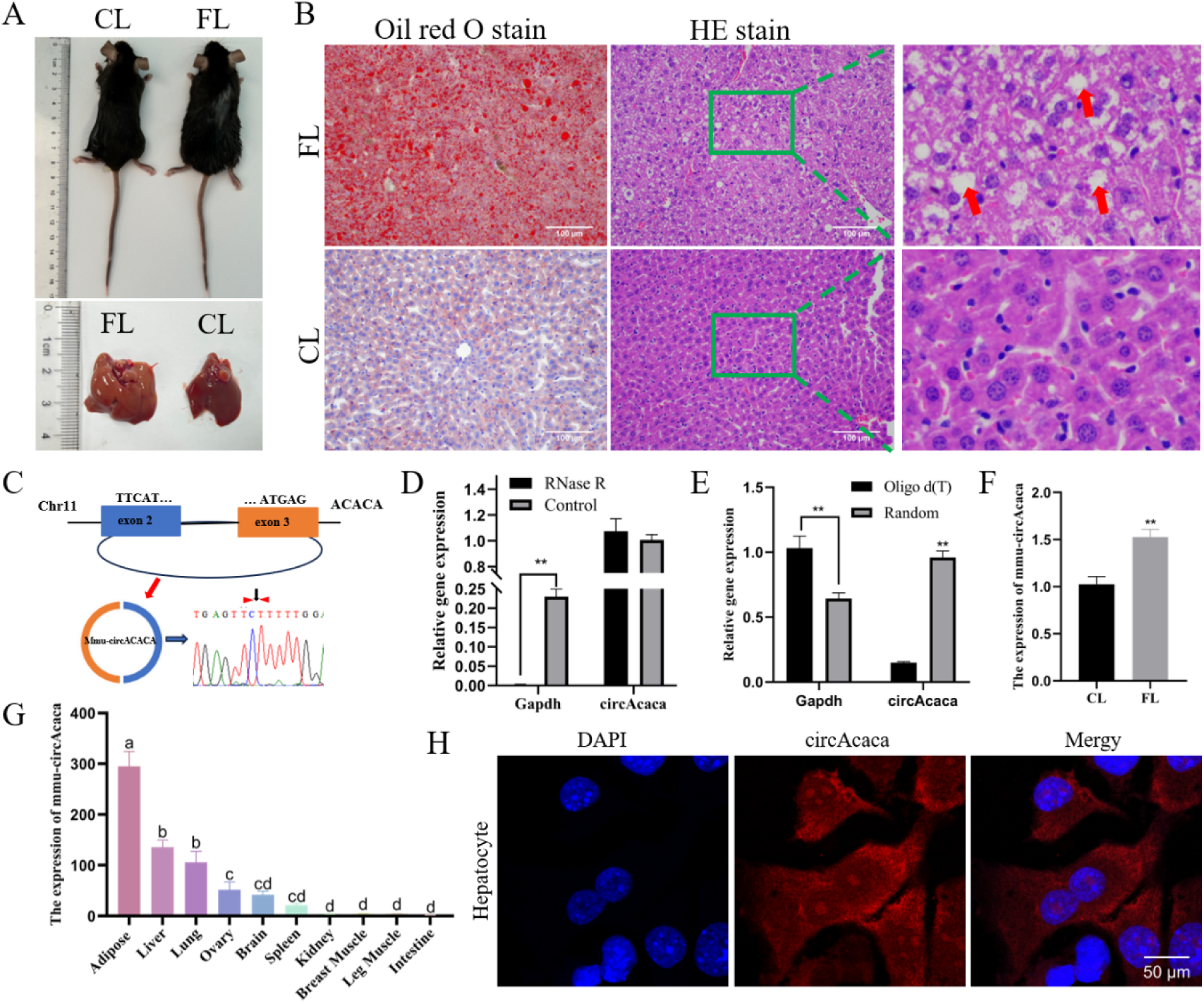
Conservation analysis of circACACA and validation of the circularity and expression pattern of mmu-circAcaca. (A) Morphologic comparison of mouse FL and CL. (B) Oil Red O staining and HE staining for FL and CL of mice. (C) Origin information of mmu- circAcaca and Sanger sequencing validates the back-splicing junctions. Black arrows show splice sites. (D) RNase R analysis of mmu-circAcaca digestion resistance. (E) Validation of the cyclic characterization of mmu-circAcaca using Oligo d(T) primers. (F) qPCR evaluation of the expression of mmu-circAcaca in CL and FL of mice. (G) qPCR analysis of the mRNA level of mmu-circAcaca in different tissues of mice. (H) FISH experiment measurement of subcellular localization of mmu-circAcaca in mouse liver cells. Scale bars= 50 μm. The data is represented as mean ± SEM. * *P* < 0.05; ** *P* < 0.01.

Analogous to chicken circACACA, mmu-circAcaca is derived from exons 2-3 of the mouse Acaca gene. Sanger sequencing confirmed the precise head-to-tail splicing sites of mmu-circAcaca (Fig 10C). Furthermore, a RNase R digestion resistance assay and amplification using an Oligo d(T) primer provided additional evidence for the existence and stability of mmu-circAcaca (Fig 10D and 10E). Notably, mmu-circAcaca was found to be highly expressed in mouse FL tissue compared to normal liver tissue (Fig 10F). Additionally, mmu-circAcaca exhibited preferential enrichment in adipose, liver, and lung tissues of the mouse (Fig 10G), and it is localized in both the nucleus and cytoplasm, with a predominant residence in the cytoplasm (Fig 10H). Collectively, these findings indicate that the structure of circACACA is highly conserved and that it stably exists in mouse liver tissues.

### Validation of the functional conservation of mmu-circAcaca in mouse liver cells

We designed and synthesized specific siRNA and overexpression vectors targeting the sequence of mmu-circAcaca to modulate its expression in vitro, with the objective of investigating its impact on mouse liver health (S10C and S10D Fig). Consistent with the effects observed for gga-circACACA, mmu-circAcaca promoted lipid accumulation in mouse hepatocytes (S10E and S10F Fig), while simultaneously impairing lipid metabolism and transport capabilities (S10G-S10J Fig). Reduction of mmu-circAcaca expression led to a significant decrease in FASN protein levels and an increase in ACOX1 (Fig 11A; S10K Fig). Conversely, overexpression of mmu- circAcaca resulted in the opposite effects (Fig 11B; S10L Fig). The group transfected with si-circAcaca exhibited reduced levels of TG and TC, whereas overexpression of mmu-circAcaca notably enhanced their accumulation (Fig 11C and 11D). Furthermore, BODIPY 493/503 staining revealed a substantial accumulation of lipid droplets within cells following mmu-circAcaca overexpression (S10M and S10N Fig). In conclusion, these findings suggest that mmu-circAcaca disrupts the balance of lipid metabolism in mouse liver cells.

**Figure. 11.**
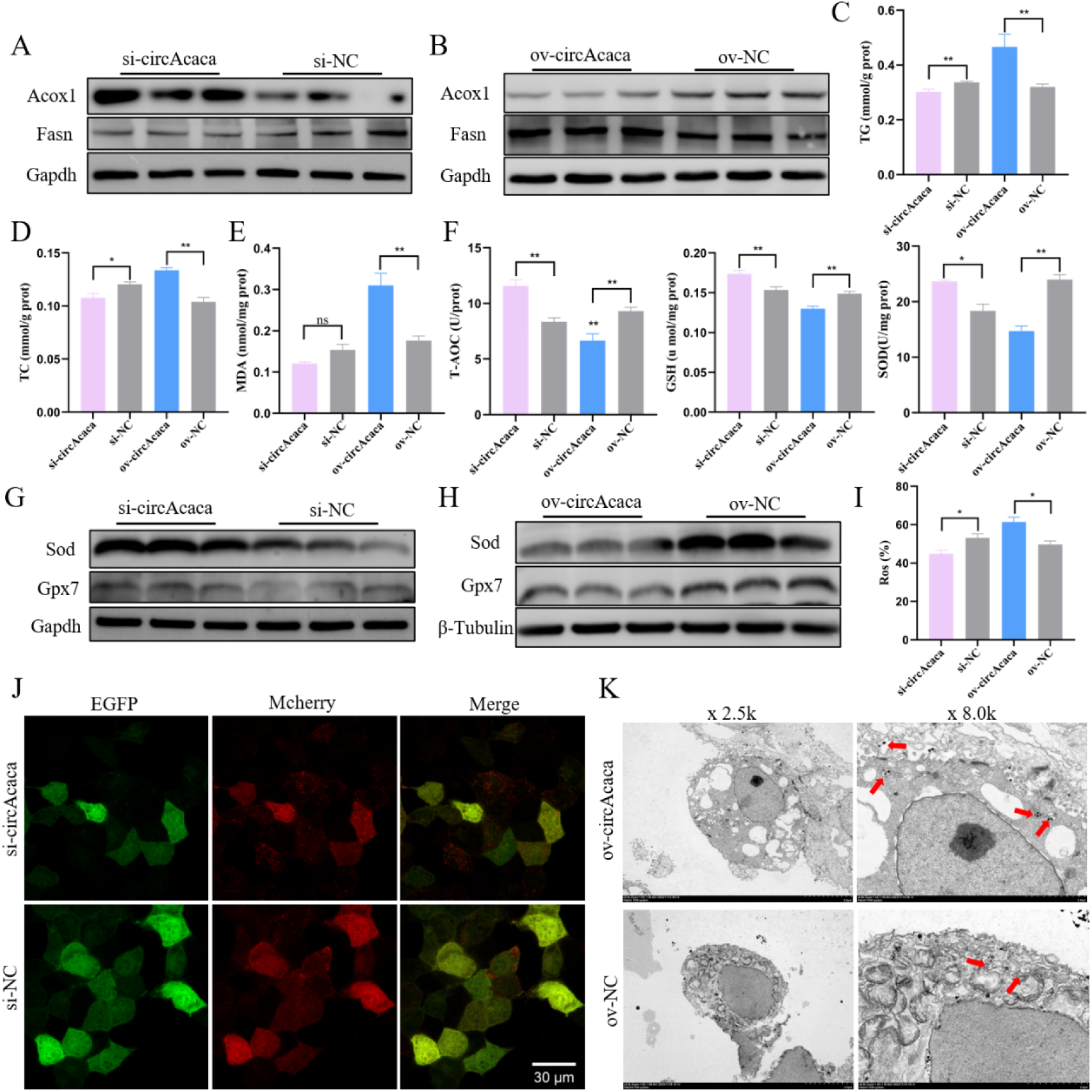
Validation of the functional conservation of mmu-circAcaca in mouse liver cells. (A, B) Western blot assessment of the protein expression of Acox1 and Fasn after silencing and overexpression of mmu-circAcaca. (C, D) Assessment of cellular TG and TC concentrations following modulation of mmu-circAcaca expression. (E, F) Quantification of the activity of cellular MDA and anti-oxidative stress related enzyme after adjusting mmu-circAcaca expression. (G, H) Western blot analysis of the protein expression of Sod and Gpx7 after the interference and overexpression of mmu-circAcaca. (I) Quantification of intracellular ROS proportions after modifying mmu-circAcaca expression. (J) Fluorescence image of Mcherry-EGFP-LC3 adenovirus construct following mmu-circAcaca inhibition. Scale bars= 30 μm. (K) Transmission electron micrograph of mice liver cells following mmu-circAcaca overexpression. The data is represented as mean ± SEM. * *P* < 0.05; ** *P* < 0.01.

Further, we found that mmu-circAcaca promoted the production of MDA (Fig 11E), while reducing the levels of antioxidant enzymes, including T-AOC, GSH, and SOD (Fig 11F). And knockdown of circAcaca increased the mRNA levels of *Gst*, *Gpx7* and *Sod*, whereas their expression was downregulated upon overexpression of mmu-circAcaca (S11A and S11B Fig). Consistent with the changes observed at the mRNA level, the protein content of oxidative stress-related genes was also fluctuated following the alteration of mmu-circAcaca expression (Fig 11G and 11H; S11C and S11D Fig). Flow cytometric analysis revealed a notable accumulation of ROS in cells of the ov- circAcaca group. Conversely, a reduction in mmu-circAcaca expression was accompanied by a decrease in intracellular ROS levels (Fig 11I; S11E Fig). In summary, these findings indicate that mmu-circAcaca impairs the antioxidant capacity of mouse liver cells.

After interfering with mmu-circAcaca, the mRNA expression of autophagy-related genes was significantly suppressed, with the exception of *P62* (S11F Fig). Conversely, overexpression of mmu-circAcaca had the opposite effect (S11G Fig). Consistent with these findings, the protein expression of autophagy-related genes also exhibited corresponding changes following the knockdown or overexpression of mmu- circAcaca (S11H and S11I Fig). Notably, the si-mmu-circAcaca treatment group demonstrated a marked reduction in the area of autophagosomes, as assessed by the Mcherry-EGFP-LC3 staining assay (Fig 12J). Furthermore, electron microscopy revealed a substantial accumulation of autolysosomes in cells overexpressing mmu- circAcaca (Fig 11K). In conclusion, similar to gga-circACACA, mmu-circAcaca also promotes autophagy in mouse hepatocytes.

**Figure. 12.**
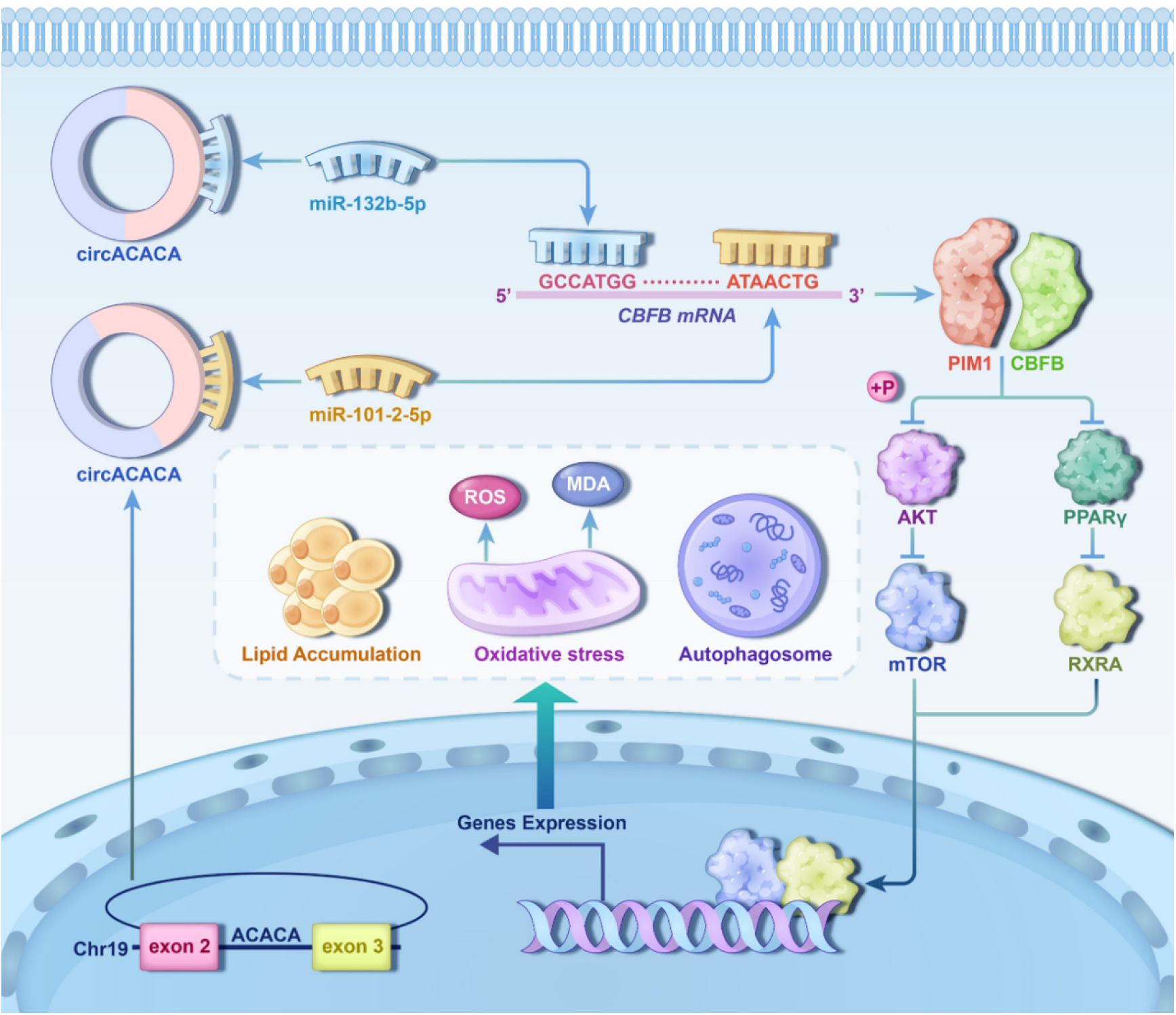
**The schematic diagram of circACACA disrupts the homeostasis of liver through acting as ceRNA of miR-132b-5p/miR-101-2-5p to regulate the activity of the AKT/mTOR and PPAR-γ signaling pathways.**

## Discussion

In recent years, there has been a notable surge in the incidence of NAFLD across all age demographics, primarily attributed to shifts in human dietary habits and lifestyles [2]. NAFLD poses a significant threat to individual liver health and presents a substantial challenge to public health systems [3, 24]. Researchers have advanced numerous hypotheses to elucidate the pathogenesis of NAFLD, with the “multiple hit” theory emerging as the most widely recognized and pertinent framework for deciphering the mechanisms driving NAFLD-associated lipotoxicity. This theory contends that disruptions in the delicate balance of hepatic fat metabolism led to an abnormal accumulation of fat within the liver. Such perturbations further disrupt the oxidative and antioxidative homeostasis of hepatocytes, culminating in the excessive generation of ROS and subsequent induction of oxidative stress. This sequence of events triggers a cascade of additional pathological processes, encompassing hepatocellular autophagy, apoptosis, inflammation, and fibrosis, thereby exacerbating overall organ damage [25, 26]. Presently, the investigation of the underlying molecular mechanisms of NAFLD is still in its early stages, and effective genetic therapeutic strategies remain elusive.

Since most circRNA sequences are highly conserved, focusing on the circRNAs that are conserved across species in animal sequence profiles may allow further opportunities for conducting clinical trials from bench to bedside [27, 28]. The genomic sequence of chickens exhibits approximately 70% homology with the human genome, and similarly, the liver, rather than adipose tissue, serves as the primary site for de novo lipid synthesis in both species [29]. Consequently, chickens have become a widely recognized model for exploring the biology of adipose tissue, metabolism, and obesity [30]. Given this context, the selection of conserved circRNAs from the chicken sequence profile is crucial for elucidating the underlying mechanisms and regulatory functions of circRNAs through cell lines and animal experiments, leveraging their shared cross-species characteristics in nucleotide sequences, biological synthesis, and functionalities [17].

FLS frequently occurs in farming poultry, particularly in laying hens. Affected chickens typically show slow growth, poor meat quality, and a marked decrease in egg production rates. In severe cases, it can lead to sudden death due to liver rupture, causing significant economic losses to the poultry industry [8, 31]. Therefore, circRNA sequencing was performed in this experiment using the chicken as a model. Through RNA sequencing and qPCR analysis, we identified several dozen circRNAs exhibiting significant differential expression between normal chicken liver and fatty liver. Among these, circACACA emerged as a particularly noteworthy candidate due to its parent gene, acetyl-CoA carboxylases alpha (ACACA), which is a key rate-limiting enzyme in de novo, fatty acid synthesis and plays a crucial regulatory role in fatty acid biosynthesis [32]. Chen et al. demonstrated that the incorporation of mulberry leaves into the diet of laying hens at a level of 3% effectively mitigates hepatic lipid accumulation by downregulating the expression of circACACA [33]. In addition, circACC1 derived from ACC1 (encoded by ACACA) pre-mRNA exerts its regulatory function on fatty acid β-oxidation and glycolysis by modulating the assembly and activation of the AMPK complex in human [34]. Consequently, we selected circACACA for subsequent validation experiments among the numerous differentially expressed circRNAs, A high-fat diet promotes hepatic lipid accumulation, inducing liver injury and oxidative stress, while impairing hepatic antioxidant defense mechanisms, modulating the expression of antioxidant-related signaling molecules, and ultimately activating autophagic pathways in the liver [35]. CircRNA plays a crucial regulatory role in this process. For example, circSCAR mitigated high-fat diet induced cirrhosis and insulin resistance in mice[36]; circPI4KB promoted intrahepatic lipid accumulation in NAFLD by mediating the expression of miR-122 [37]. CircCBFB facilitated acetaminophen (APAP)-induced liver injury and mitochondrial oxidative stress through the modulation of p66Shc expression [38]; mmu_circ_0009303 enhanced oxidative stress, inflammation, and excessive fat accumulation in liver cells induced by free fatty acids (FFA) through modulating the miRNA-182-5p/Foxo3 pathway and lipid metabolism-related regulatory proteins [39]. Overexpression of circLDLR significantly reduced the accumulation of lipid droplets, lowered the levels of TG, TC, and p62, and increased the levels of LC3-II along with the number of autophagosomes in the NAFLD cell model induced by oleic acid and palmitic acid [40]; circRNA_002581 knockdown accelerated the degree of autophagy and dramatically decreased the generation of lipid droplets in NASH cell and mouse models [41]. Our study also revealed that circACACA contributes to a notable accumulation of fat, oxidative stress and autophagy in the chicken liver.

CircRNAs typically function as ceRNAs, containing miRNA response elements, to bind to miRNAs and alleviate their inhibitory effects on target genes, thereby exerting their functional mechanisms. Previous studies have demonstrated that circACACA regulated occurrence and development of multiple tumors through the ceRNA mechanism. For example, circACACA facilitated colorectal tumorigenesis by regulating the miR-193a/b-3p/HDAC3/p53 axis to activate the MVA pathway [42]; circACACA enhanced migration, invasion, proliferation and glycolysis of cervical cancer cells by acting as sponges of miR-582-5p [43]; circACACA impaired the cellular functions of non-small cell lung cancer cells, including proliferation, invasion, and migration, via regulation of the miR-1183 and PI3K/AKT signaling pathways [44]. Although circACACA has been implicated in hepatic lipid metabolism, its molecular regulatory mechanism remains unreported. In this study, we found that circACACA can simultaneously target miR-132b-5p and miR-101-2-5p to regulate the formation of FL in chicken. Previous studies have also found a close association between them and liver diseases. Halin et al. established miR-132 as a critical regulator of hepatic lipid homeostasis, as evidenced by the severe hepatic steatosis, obesity, hyperlipidemia (elevated serum LDL/VLDL), and TG accumulation observed in miR-132 overexpressing transgenic mice [45]. Xu et al. reported that autophagy was suppressed by miR-101, and enhanced cisplatin-induced apoptosis in hepatocellular carcinoma cells [46]. Our research findings further revealed that miR-132b-5p and miR-101-2-5p are intricately intertwined with lipid metabolism, oxidative stress, and autophagic processes within the liver.

Subsequent bioinformatics predictions combined with experimental validation identified the core-binding factor subunit beta (CBFB) gene as a shared target of both miR-132b-5p and miR-101-2-5p. CBFB, recognized as a transcriptional co-factor for RUNX family proteins, plays a critical role in enhancing DNA binding capacity of these complexes [47]. This stabilization mechanism fundamentally regulates gene expression by promoting target gene transcription [48]. While CBFB is well-established for its functions in gene regulation, cellular differentiation, hematopoiesis, and bone development, its regulatory roles in metabolic processes remain under investigation. Our study elucidates CBFB’s novel regulatory capacity in controlling hepatic fat metabolism, oxidative stress responses, and autophagic processes. Mechanistic investigations revealed that CBFB inhibits AKT activation through downregulation of protein kinase PIM1 [49]. As a member of the serine/threonine kinase superfamily, PIM1 modulates multiple signaling pathways through distinct molecular interactions [50]. Notably, PIM1’s interaction with mTOR complex 1 (mTORC1) establishes a pivotal regulatory axis for lipid metabolism and protein synthesis [51, 52]. Consistent with previous findings, Zhang et al. demonstrated that PTUPB induction promotes autophagy, attenuates hepatocyte senescence, and ameliorates NAFLD progression through PI3K/AKT/mTOR pathway inhibition [53]. Parallel studies have established that AKT/mTOR signaling exerts dual regulatory effects in liver steatosis management promoting oxidative stress resolution while enhancing autophagic flux [54, 55]. Our experimental validation confirmed the functional interaction between CBFB and PIM1, revealing that circACACA exerts its metabolic effects through indirect inactivation of the AKT/mTOR pathway via modulation of the CBFB/PIM1 complex. Furthermore, PIM1 was vital for the metabolism of fat via mediating PPAR-γ signal pathway [56, 57]. PPAR-γ, as a master regulator of hepatic lipid homeostasis, demonstrates therapeutic potential for NAFLD treatment through its anti-inflammatory, insulin-sensitizing, oxidative stress-alleviating, and fibrotic regression properties [58]. Interestingly, our study uncovered a novel inhibitory effect of circACACA on PPAR-γ signaling through the same CBFB/PIM1 axis. In summary, this comprehensive investigation demonstrates that circACACA reduces the activity of the AKT/mTOR and PPAR-γ signaling pathways by targeting miR-132b-5p/miR-101-2-5p and CBFB/PIM1 axis, thereby modulating liver metabolism and healthy.

Evolutionary biology has demonstrated that essential genetic elements critical for organismal survival and reproductive success exhibit remarkable stability under environmental pressures and selective constraints [59, 60]. This evolutionary stability not only sustains biological diversity but also provides foundational mechanisms for speciation. Notably, circRNAs represent evolutionarily conserved regulatory molecules that play integral roles in gene expression networks [27]. Comparative genomics studies across species have revealed substantial sequence conservation among circRNAs, underscoring their fundamental biological importance [28]. Liang et al.’s work elucidated key molecular characteristics, conserved sequences, and spatiotemporal expression patterns of identified 5,934 circRNAs across nine organs and three skeletal muscles in pigs, demonstrating significant homology with human and murine counterparts [61]. For instance, circCDR1as (also known as ciRS-7) exhibits striking sequence conservation across mammals, with conserved ceRNA functionality for miR- 7 in both humans and mice, regulating critical processes such as neurodevelopment and oncogenesis [62–64]. Similarly, circZNF609 displays conserved expression patterns and muscle-specific functions in humans and mice [65], while circMEF2As demonstrate evolutionary conservation in promoting muscle development across chickens and rodents [66]. To investigate circACACA’s evolutionary trajectory, we utilized the circAtlas database for comparative sequence analysis across human, murine, and avian species. Our findings revealed 72.67% sequence similarity between chicken and human circACACA, and 69.67% homology with murine orthologs. Notably, circACACA originates from exons 2 and 3 of the ACACA gene in all examined species, maintaining a consistent 300-nucleotide length. This conservation pattern suggests potential functional relevance across vertebrates. To experimentally validate these bioinformatic predictions, we established a murine model system to characterize circAcaca’s structural integrity and functional conservation. Through circular RNA validation assays, we confirmed its stable circular structure in hepatic tissues. Functional studies further demonstrated conserved roles in regulating lipid metabolism, oxidative stress responses, and autophagic flux in mice.

Currently, circRNA has garnered extensive attention in the field of human medicine.

It has been well-established that numerous circRNAs contribute to the pathogenesis of malignancies and are regarded as potential biomarkers for disease diagnosis and prognosis [67, 68]. Moreover, owing to their slow degradation rate and stable presence in the body, circRNAs can exert their functional roles by interacting with miRNAs or producing substantial amounts of proteins. Consequently, they are viewed as a novel therapeutic approach for human diseases and have emerged as promising targets for drug development [14, 69]. In our study, we utilized chickens and mice as research models to comprehensively investigate the functions and mechanisms of circACACA in liver fat metabolism and related diseases, as well as to thoroughly evaluate its potential influence on the progression of NAFLD. Our findings indicate that circACACA may serve as a potential biomarker for NAFLD.

## Conclusion

This study identifies circACACA as a novel FL-enriched circRNA that functions as a ceRNA to disrupt the miR-132b-5p/miR-101-2-5p-CBFB regulatory axis, thereby repressing PIM1 expression and attenuating AKT/mTOR and PPAR-γ signaling, which collectively perturb hepatic lipid metabolism, exacerbate oxidative stress, induce autophagy, and ultimately impair liver function (Fig 12). Notably, its conserved regulation across species highlights its pivotal role in hepatic metabolism, which suggests promising translational applications for both enhancing livestock production traits and developing targeted therapies for NAFLD.

## Acknowledgments

We would like to express our sincere gratitude to our supervisors, Prof. Qing Zhu and Prof. Huadong Yin, for their invaluable guidance and insightful feedback throughout this project. We thank Dr. Shunshun Han for his technical guidance throughout the trial. Finally, I would like to thank my university, Sichuan Agricultural University, for providing us with a better research platform.

## Author Contributions

Conceptualization: Shunshun Han, Huadong Yin, Qing Zhu; Formal analysis: Jing Zhao, Shunshun Han, Jialin Xiang; Methodology: Jing Zhao, Yuqi Chen, Xiyu Zhao, Chang Liu; Funding acquisition: Shunshun Han, Qing Zhu, Huadong Yin; Validation: Jing Zhao, Jialin Xiang, Wenjuan Wang; Data curation: Jing Zhao, Shunshun Han, Yao Zhang; Writing-original draft: Jing Zhao, Shunshun Han; Writing-review & editing: Qing Zhu, Chang Liu, Huadong Yin;

## Data Availability Statement

All relevant data are submitted with the manuscript as source data files. CircRNA sequencing raw data have been submitted to the SRA database at NCB1 under the accession number PRJNA960781.

## Supporting information

**S1 Fig.**
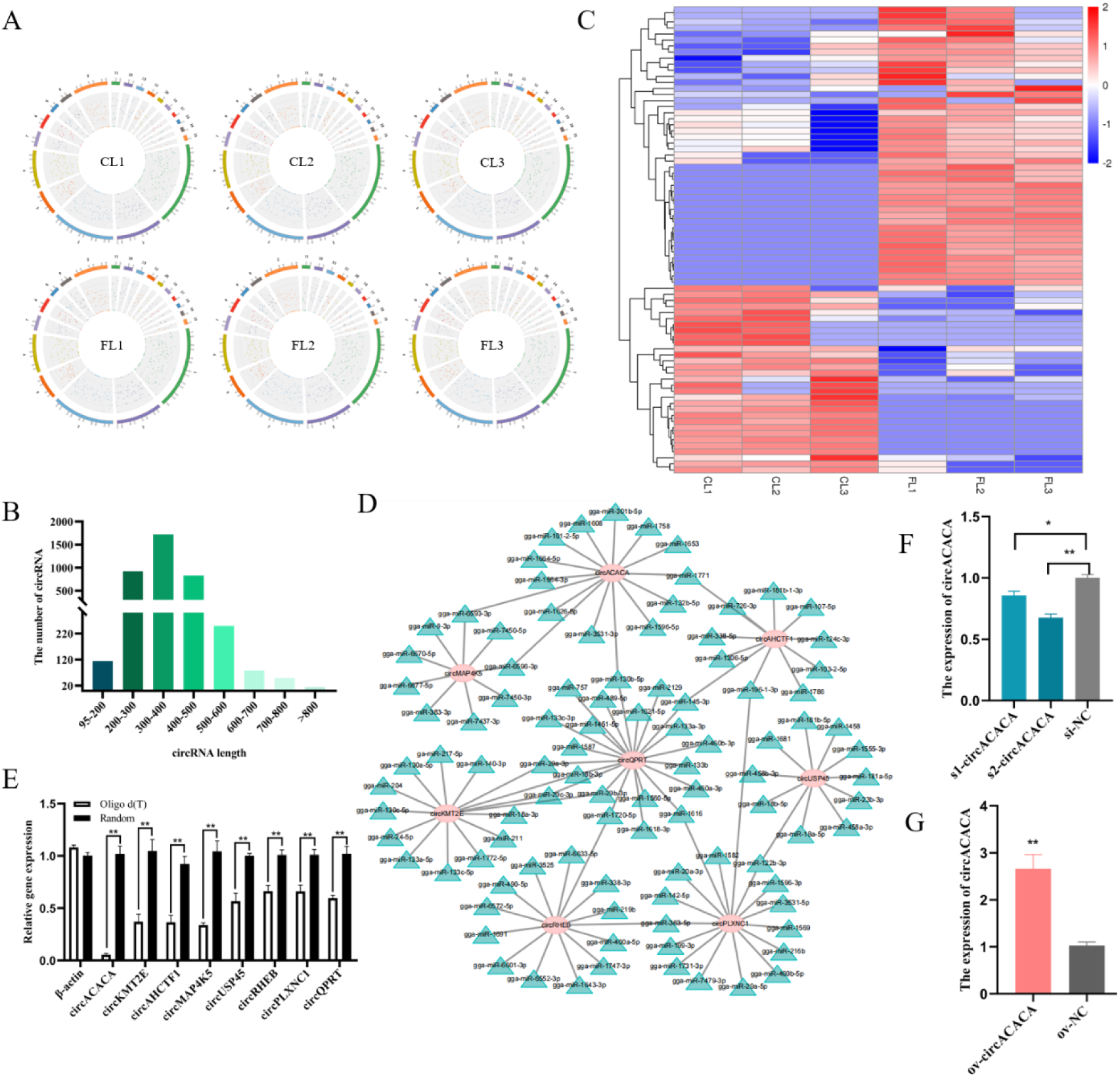
Overview of circRNA deep sequencing data and analysis of differentially expressed circRNAs. (A) Distribution density of reads from all samples on the chicken chromosomes. (B) Sequence length statistics of circRNAs. (C) Cluster heat map of differentially expressed circRNAs in the CL and FL. (D) Network diagram of the top 8 differentially high-expressing circRNAs and their target genes. (E) qPCR evaluation of the top 8 differentially high-expressing circRNAs and β-actin expression using oligo d(T) and random primers form chicken liver. (F) qPCR analysis of the expression of circACACA after transfected small interference RNA (s1-circACACA, s2-circACACA and si-NC). (G) qPCR assessment of the expression of circACACA after transfected overexpression plasmid (ov- circACACA and ov-NC). The data is represented as mean ± SEM. * *P* < 0.05; ** *P* < 0.01.

**S2 Fig.**
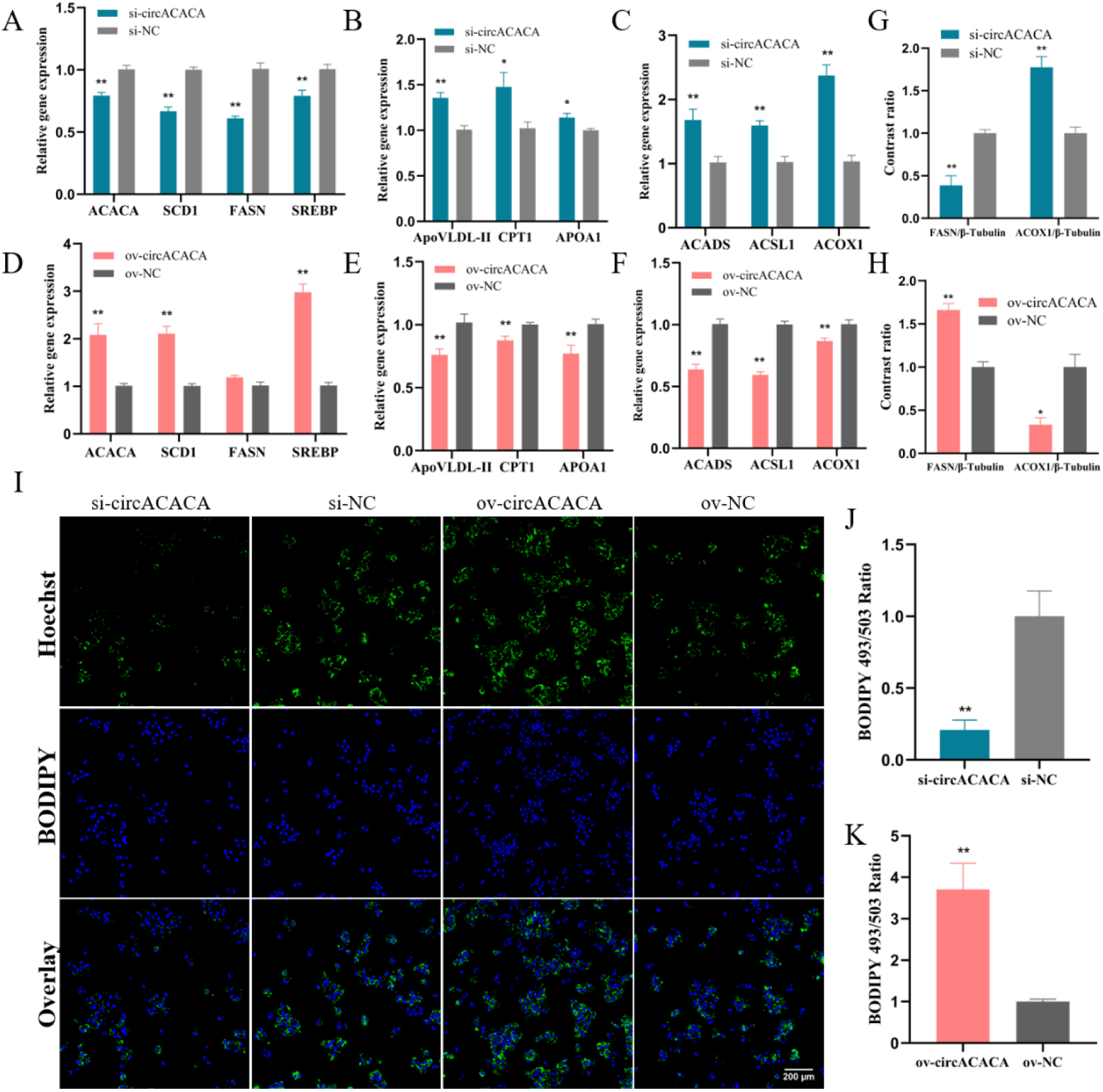
circACACA damages liver healthy by promoting lipid accumulation, oxidative stress and autophagy in chicken hepatocytes. (A) qPCR analysis of the mRNA expression of fat synthesis genes (*ACACA*, *SCD1*, *FASN*, and *SREBP*) after circACACA interference. (B) qPCR analysis of the mRNA level of lipid transport-related genes (ApoVLDL-Ⅱ, APOA1 and CPT1) after circACACA interference. (C) qPCR evaluation of the expression of β-oxidation related genes (ACSL1, ACADS and ACOX1) after circACACA interference. (D-F) qPCR evaluation of the expression fat synthesis genes, lipid transport-related genes and β-oxidation related genes after increasing of circACACA. (G, H) Statistics analysis of gray values for the FSAN and ACOX1 protein after knockdown and overexpression of circACACA. (I) BODIPY 493/503 staining assessment of lipid droplets after changing the expression of circACACA. Scale bars= 200 μm. (J, K) Statistics on the area of positive staining for BODIPY 493/503 staining after interference and overexpression of circACACA. The data is represented as mean ± SEM. * *P* < 0.05; ** *P* < 0.01.

**S3 Fig.**
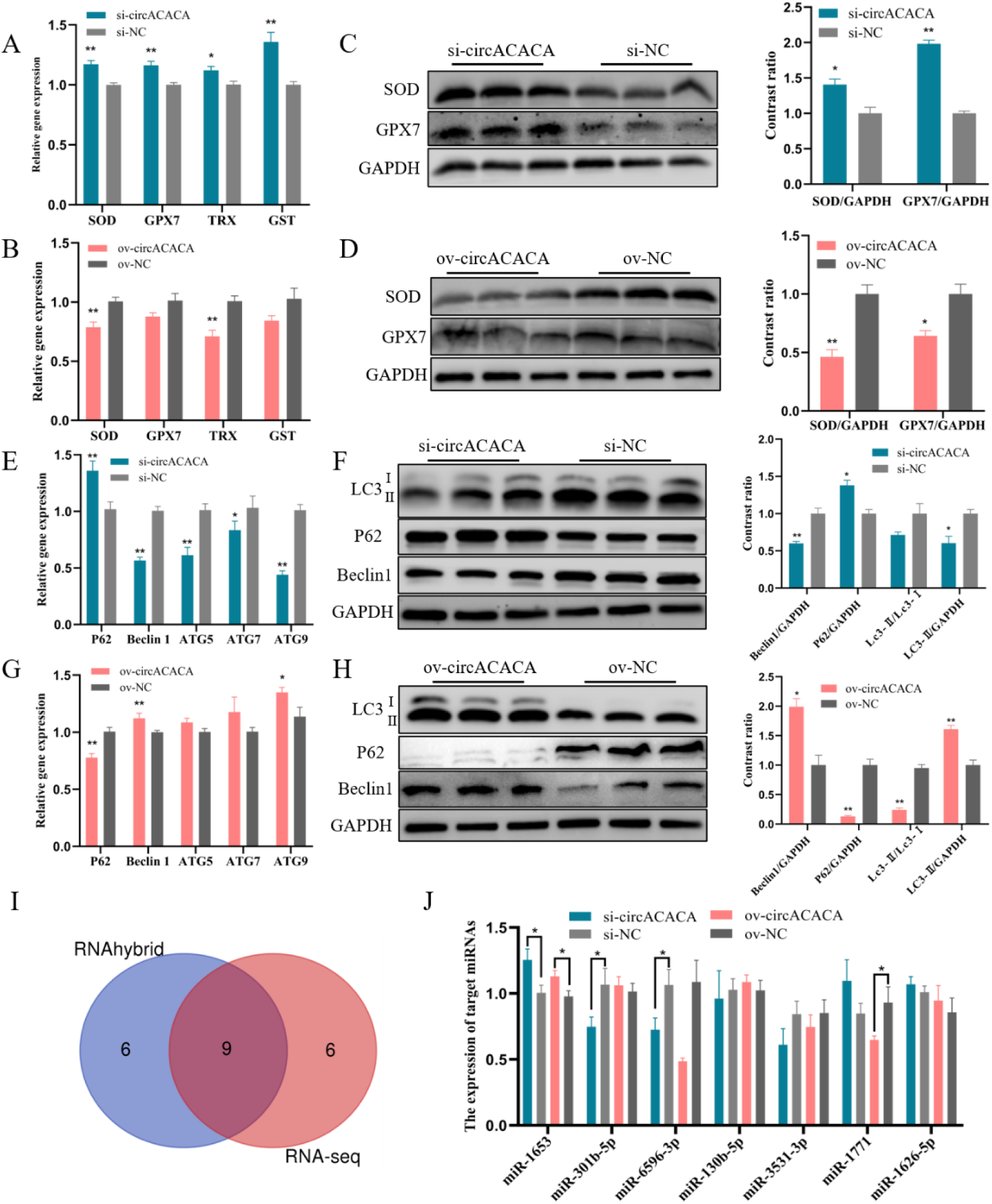
circACACA damages liver healthy by promoting lipid accumulation, oxidative stress and autophagy in chicken hepatocytes. (A, B) qPCR analysis of the mRNA level of anti-oxidative stress related gene (SOD, GPX, TRX, and GST) after interference and overexpression circACACA. (C, D) Western blot analysis of the protein level of SOD and GPX7 after decreasing and increasing of circACACA. (E) qPCR measurement of the mRNA expression of the autophagy related gene (P62, Beclin1, ATG5, ATG7, and ATG9) after knockdown of circACACA. (F) Western blot evaluation of the protein level of LC3, P62 and Beclin1 after circACACA interference. (G) qPCR analysis of the mRNA level the P62, Beclin1, ATG5, ATG7, and ATG9 after overexpression of circACACA. (H) Western blot examination of the protein level of LC3, P62 and Beclin1 after addition of ov-circACACA. (I) Venn diagram of circRNA target miRNA predictions using RNAhybrid and circRNA-seq. (J) qPCR examination of the expression of target miRNAs after interference and overexpression of circACACA. The data is represented as mean ± SEM. * *P* < 0.05; ** *P* < 0.01.

**S4 Fig.**
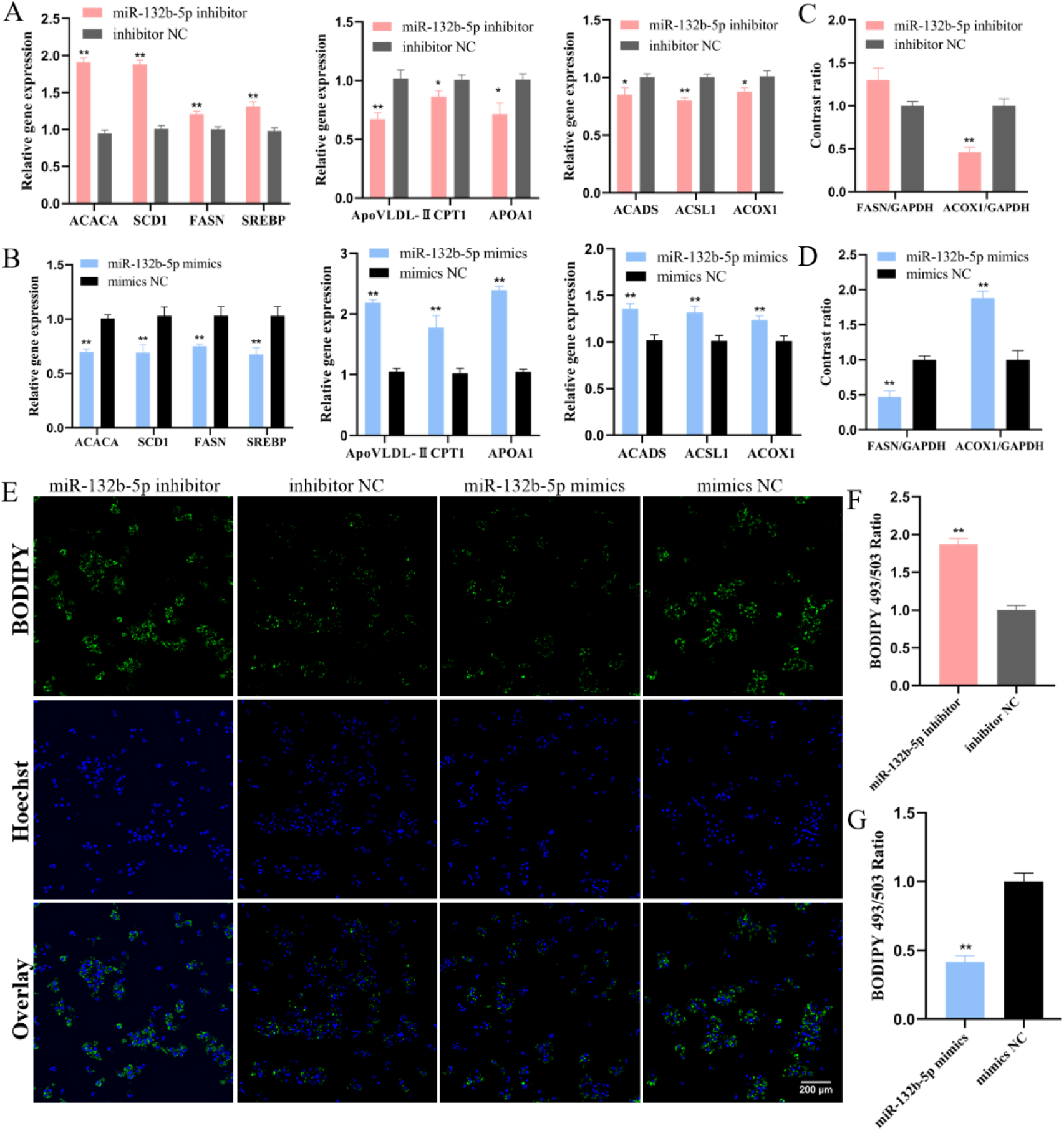
miR-132b-5p and miR-101-2-5p contribute to the health of the liver by mitigating fat accumulation, oxidative damage and autophagy in chicken hepatocytes. (A, B) qPCR evaluation of the expression of lipid metabolism related genes after inhibition and overexpression of miR-132b-5p. (C, D) Statistics of FASN and ACOX1 protein gray value after knockdown and upregulation of miR-132b-5p. (E) BODIPY 493/503 staining of lipid droplets after knockdown and overexpression of miR-132b-5p. Scale bars= 200 μ m. (F, G) The statistics of BOD 493/503 staining area after changing miR-132b-5p. The data is represented as mean ± SEM. * *P* < 0.05; ** *P* < 0.01.

**S5 Fig.**
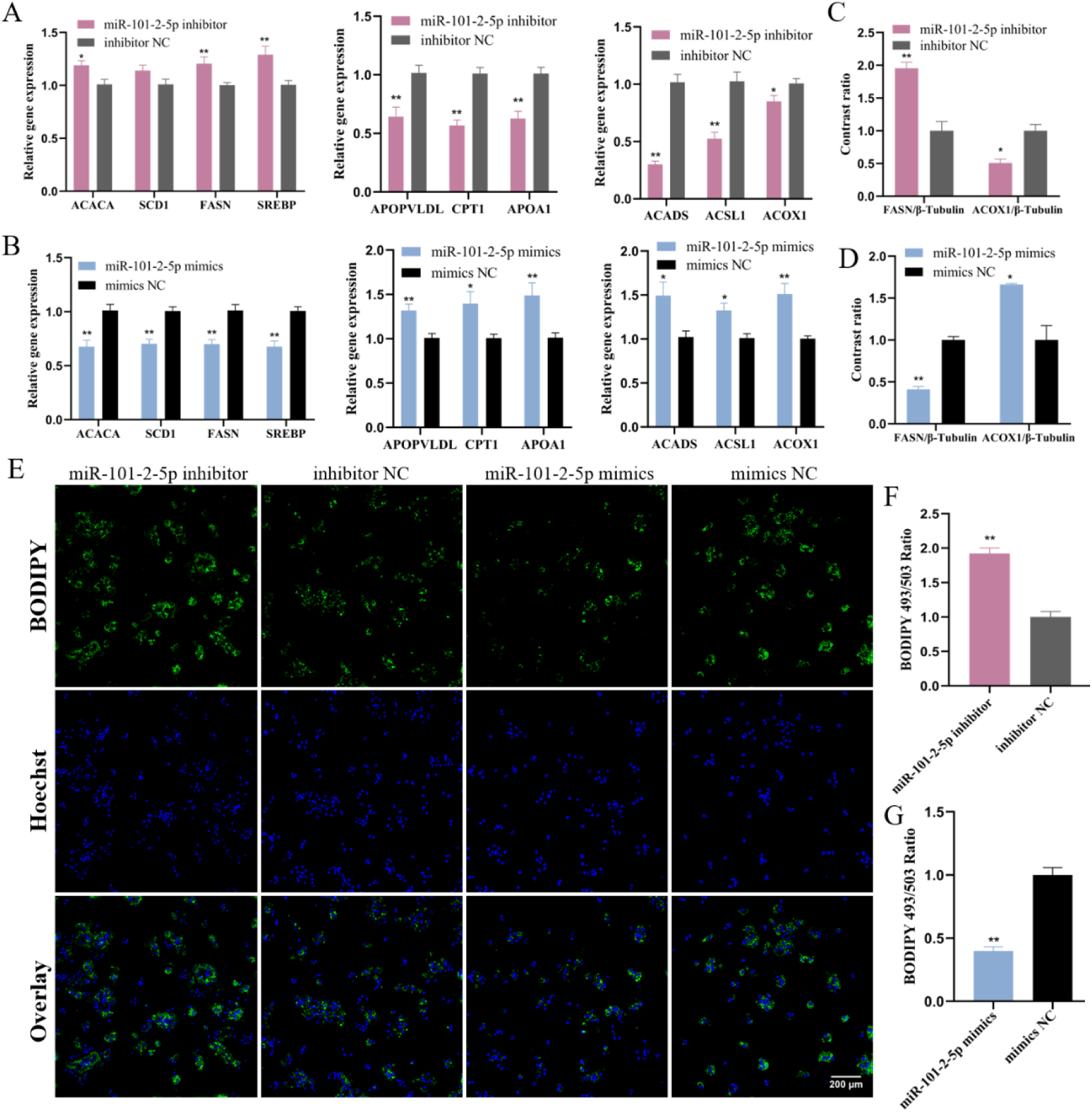
miR-132b-5p and miR-101-2-5p contribute to the health of the liver by mitigating fat accumulation, oxidative damage and autophagy in chicken hepatocytes. (A, B) qPCR evaluation of the expression of lipid metabolism related genes after interference and overexpression of miR-101-2-5p. (C, D) Statistics of FASN and ACOX1 protein gray value after interference and overexpression of miR-101-2-5p. (E) BODIPY 493/503 staining of lipid droplets after interference and upregulation of miR-101-2-5p. Scale bars= 200 μm. (F, G) The statistics of BOD 493/503 staining area after altering miR-101-2-5p. The data is represented as mean ± SEM. * *P* < 0.05; ** *P* < 0.01.

**S6 Fig.**
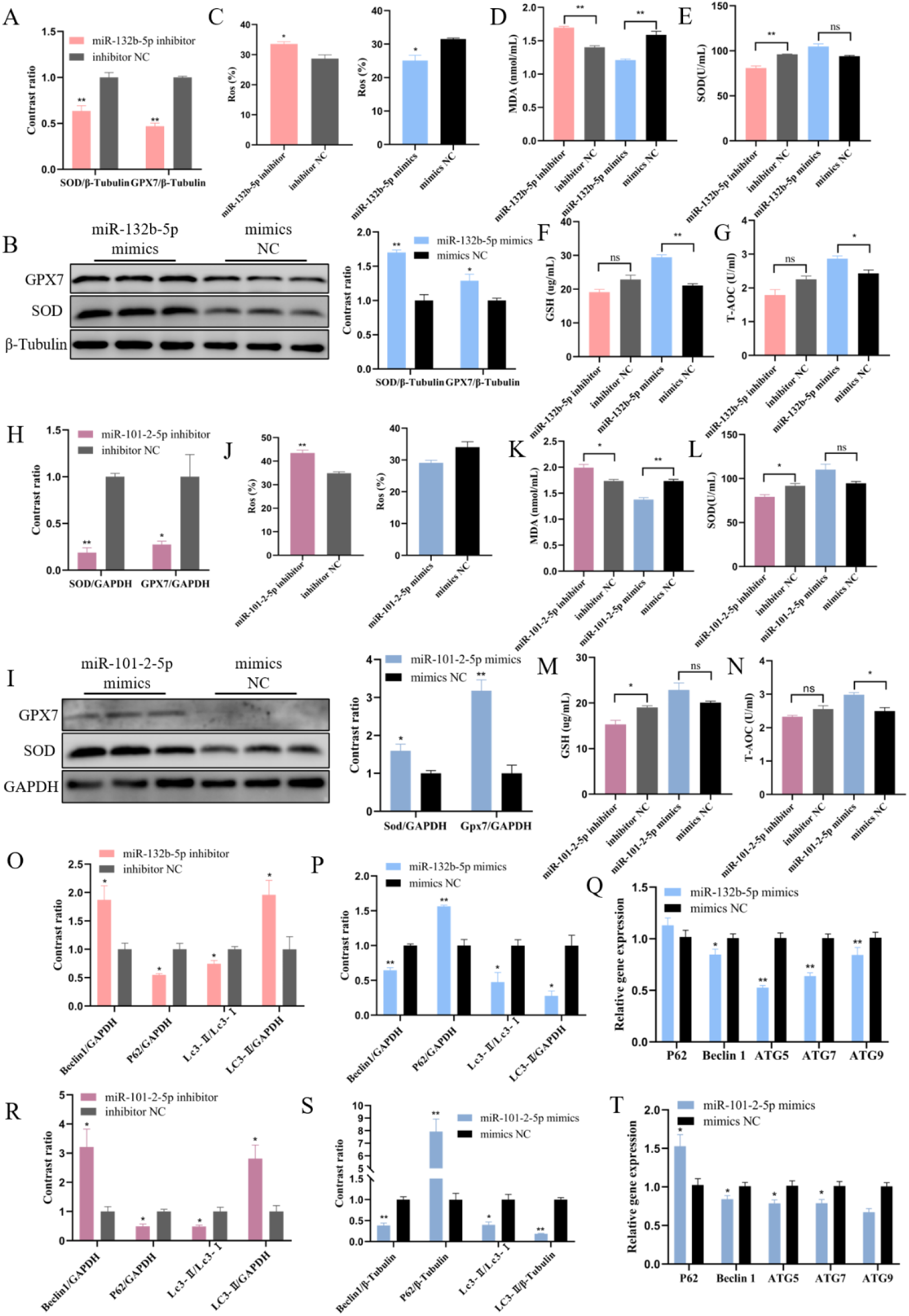
miR-132b-5p and miR-101-2-5p contribute to the health of the liver by mitigating fat accumulation, oxidative damage and autophagy in chicken hepatocytes. (A) Statistics analysis of gray values for the GPX7 and SOD protein after knockdown of miR-132b-5p. (B) Western blot analysis of the protein level of GPX7 and SOD after overexpression of miR-132b- 5p. (C) Quantification of intracellular ROS proportions after changing expression miR-132b- 5p. (D-G) The accumulation level of MDA, SOD, GSH and T-AOC after changing the expression of miR-132b-5p. (H) Statistics analysis of gray values for the GPX7 and SOD protein after knockdown of miR-101-2-5p. (I) Western blot assessment of the protein level of GPX7 and SOD after upregulation of miR-101-2-5p. (J) Quantification of intracellular ROS proportions after interference and overexpression of miR-132b-5p. (K-N) The concentration of MDA, SOD, GSH and T-AOC after changing the expression of miR-101-2-5p. (O, P) Statistics of the protein gray value of autophagy related genes after interference and overexpression of miR-132b-5p. (Q) qPCR evaluation of the mRNA level of autophagy related genes after increasing of miR-132b-5p. (R, S) Statistics of the protein gray value of autophagy related genes after knockdown and overexpression of miR-101-2-5p. (T) qPCR analysis of the mRNA level of autophagy related genes after upregulation of miR-101-2-5p. The data is represented as mean ± SEM. * *P* < 0.05; ** *P* < 0.01.

**S7 Fig.**
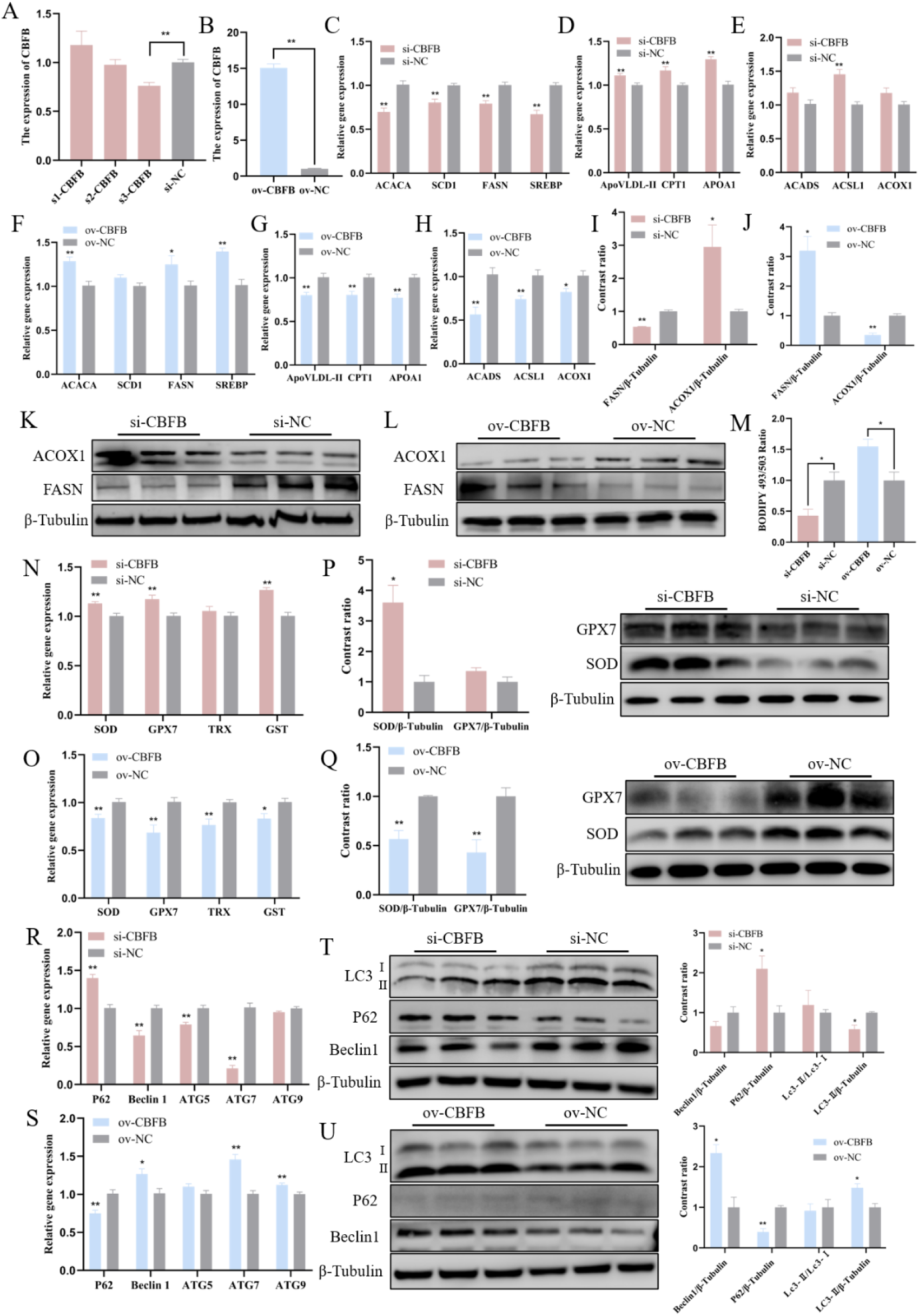
CBFB impairs liver health by promoting fat accumulation, oxidative damage and autophagy. (A) qPCR assessment of the gene expression of CBFB after different siRNA (s1- CBFB, s2-CBFB and s3-CBFB) treatments. (B) qPCR evaluation of the mRNA level of CBFB after transfection with CBFB overexpression vector (ov-CBFB). (C-E) qPCR examination of the mRNA expression of fat metabolism related gene after interference of CBFB. (F-H) qPCR analysis of the mRNA level of fat metabolism related gene after overexpression of CBFB. (I, K) Western blot measurement of the protein expression of FASN and ACOX1 following CBFB interference. (J, L) Western blot analysis of the protein expression of FASN and ACOX1 following CBFB overexpression. (M) Statistics of the BODIPY 493/503 fluorescence-stained area after altering CBFB expression. (N, O) qPCR evaluation of the mRNA level of anti- oxidative stress related gene following blocking and increasing CBFB. (P, Q) Western blot assessment of the protein level of GPX7 and SOD after interference and overexpression of CBFB. (R, S) qPCR analysis of the mRNA level of autophagy related gene following the knockdown and overexpression of CBFB. (T, U) Western blot evaluation of the protein content of autophagy related gene after interference and upregulation of CBFB. The data is represented as mean ± SEM. * *P* < 0.05; ** *P* < 0.01.

**S8 Fig.**
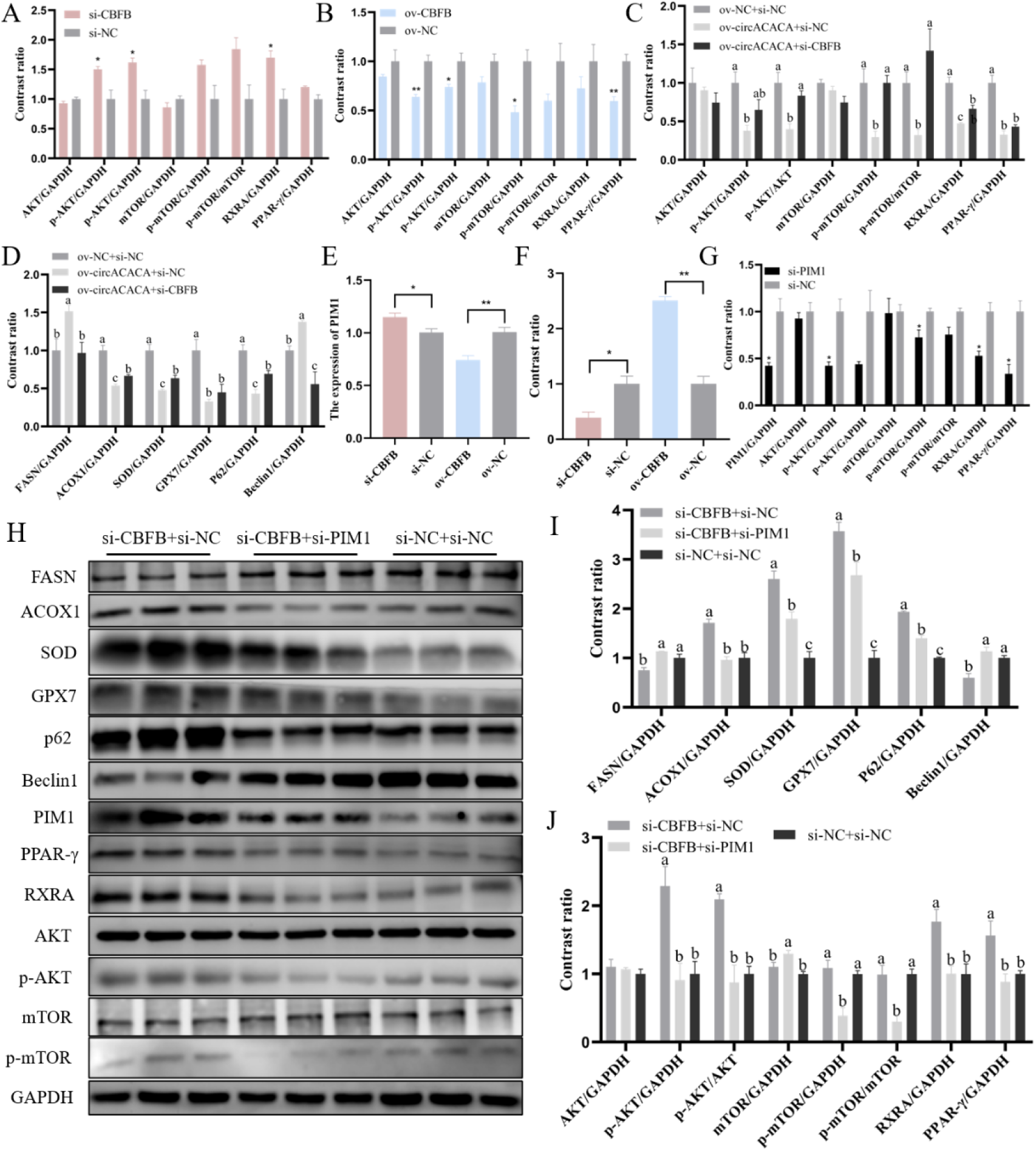
CircACACA inactivates AKT/mTOR and PPAR-. γ **signaling pathways by regulating CBFB/PIM1 complexes.** (A, B) Statistics of the gray value of key proteins in the AKT/mTOR and the PPAR-γ signaling pathway following blocking and increasing CBFB. (C) Statistics analysis of gray values for the key proteins in the AKT/mTOR and the PPAR- γ signaling pathway after co-transfected with ov-NC+si-NC and ov-circACACA+si-NC or si- CBFB. (D) Statistics analysis of gray values for fat metabolism related, anti-oxidative stress related and autophagy related proteins in co-transfection group with ov-NC+si-NC and ov- circACACA+si-NC or si-CBFB. (E) qPCR evaluation of the mRNA level of PIM1 after knockdown and overexpression of CBFB. (F) Statistics of PIM1 protein gray value after changing the expression of CBFB. (G) Statistics analysis of gray values for PIM1, p-AKT, p- mTOR, RXRA and PPAR- γ after interference of PIM1. (H) Western blot analysis of the protein levels of fat metabolism related protein, anti-oxidative stress related protein, autophagy related protein, the AKT/mTOR and the PPAR- γ signaling pathway key protein after co- transfected with si-CBFB+si-NC/si-PIM1 and si-NC+si-NC. (I) Statistics analysis of protein gray values for FASN, ACOX1, SOD, GPX7, P62 and Beclin1 after co-transfected with si- CBFB+si-NC/si-PIM1 and si-NC+si-NC. (J) Statistics analysis of protein gray values for p- AKT, p-mTOR, RXRA and PPAR-γ after co-transfected with si-CBFB+si-NC/si-PIM1 and si-NC+si-NC. The data is represented as mean ± SEM. * P < 0.05; ** P < 0.01.

**S9 Fig.**
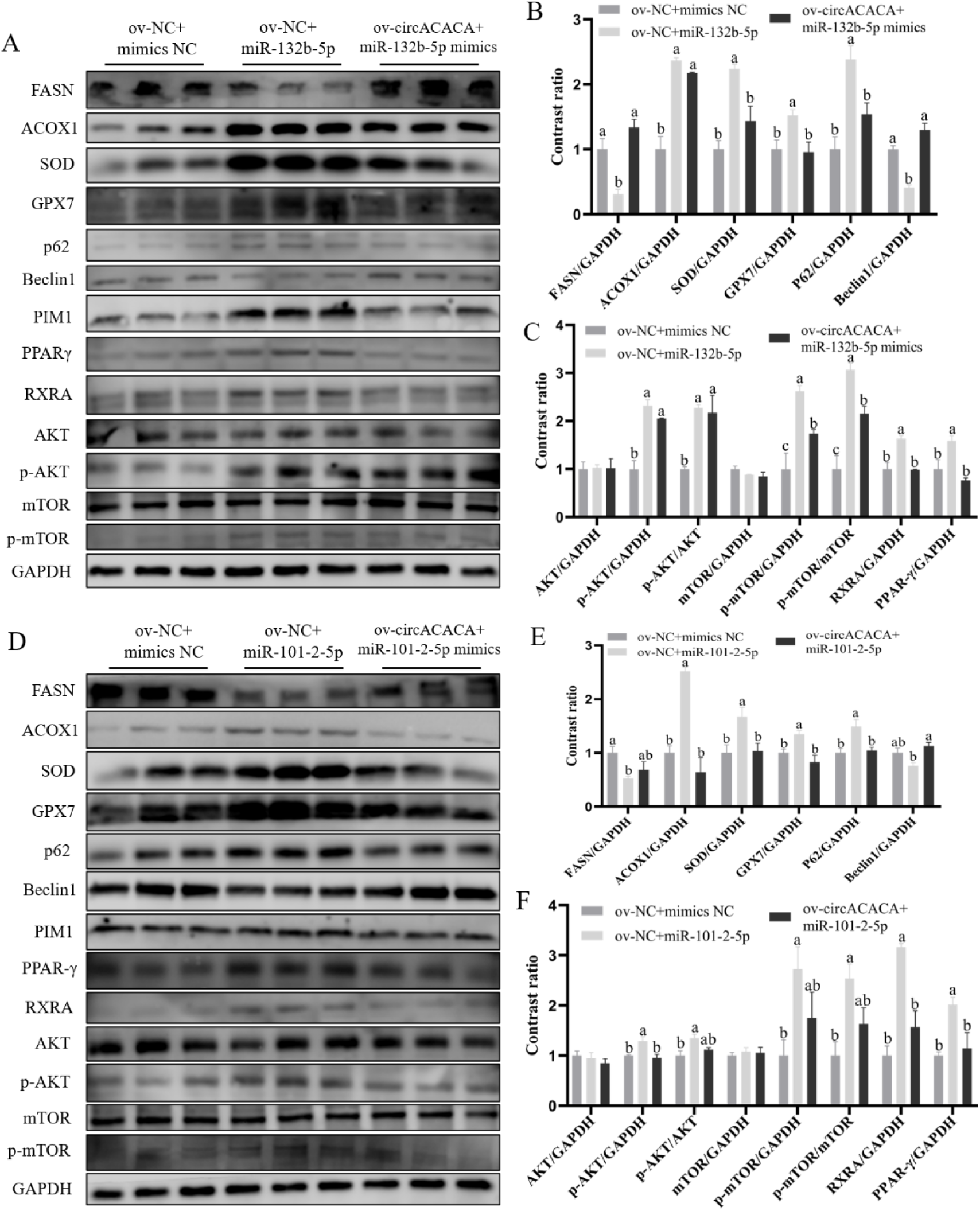
CircACACA inactivates AKT/mTOR and PPAR-. γ **signaling pathways by regulating CBFB/PIM1 complexes.** (A) Western blot analysis of the protein levels of fat metabolism related protein, anti-oxidative stress related protein, autophagy related protein, the AKT/mTOR and the PPAR- γ signaling pathway key protein after co-transfected with ov-NC+mimics NC and mimics miR-132b-5p+ov-NC or ov-circACACA. (B) Statistics analysis of protein gray values for FASN, ACOX1, SOD, GPX7, P62 and Beclin1 after co-transfected with ov-NC+mimics NC and mimics miR-132b-5p+ov-NC or ov-circACACA. (C) Statistics analysis of protein gray values for p-AKT, p-mTOR, RXRA and PPAR-γ after co-transfected with ov-NC+mimics NC and mimics miR-132b-5p+ov-NC or ov-circACACA. (D) Western blot analysis of the protein levels of fat metabolism related protein, anti-oxidative stress related protein, autophagy related protein, the AKT/mTOR and the PPAR-γ signaling pathway key protein after co-transfected with ov-NC+mimics NC and mimics miR-101-2-5p+ov-NC or ov- circACACA. (E) Statistics analysis of protein gray values for FASN, ACOX1, SOD, GPX7, P62 and Beclin1 after co-transfected with ov-NC+mimics NC and mimics miR-101-2-5p+ov- NC or ov-circACACA. (F) Statistics analysis of protein gray values for p-AKT, p-mTOR, RXRA and PPAR-γ after co-transfected with ov-NC+mimics NC and mimics miR-101-2- 5p+ov-NC or ov-circACACA. The data is represented as mean ± SEM. * *P* < 0.05; ** *P* < 0.01.

**S10 Fig.**
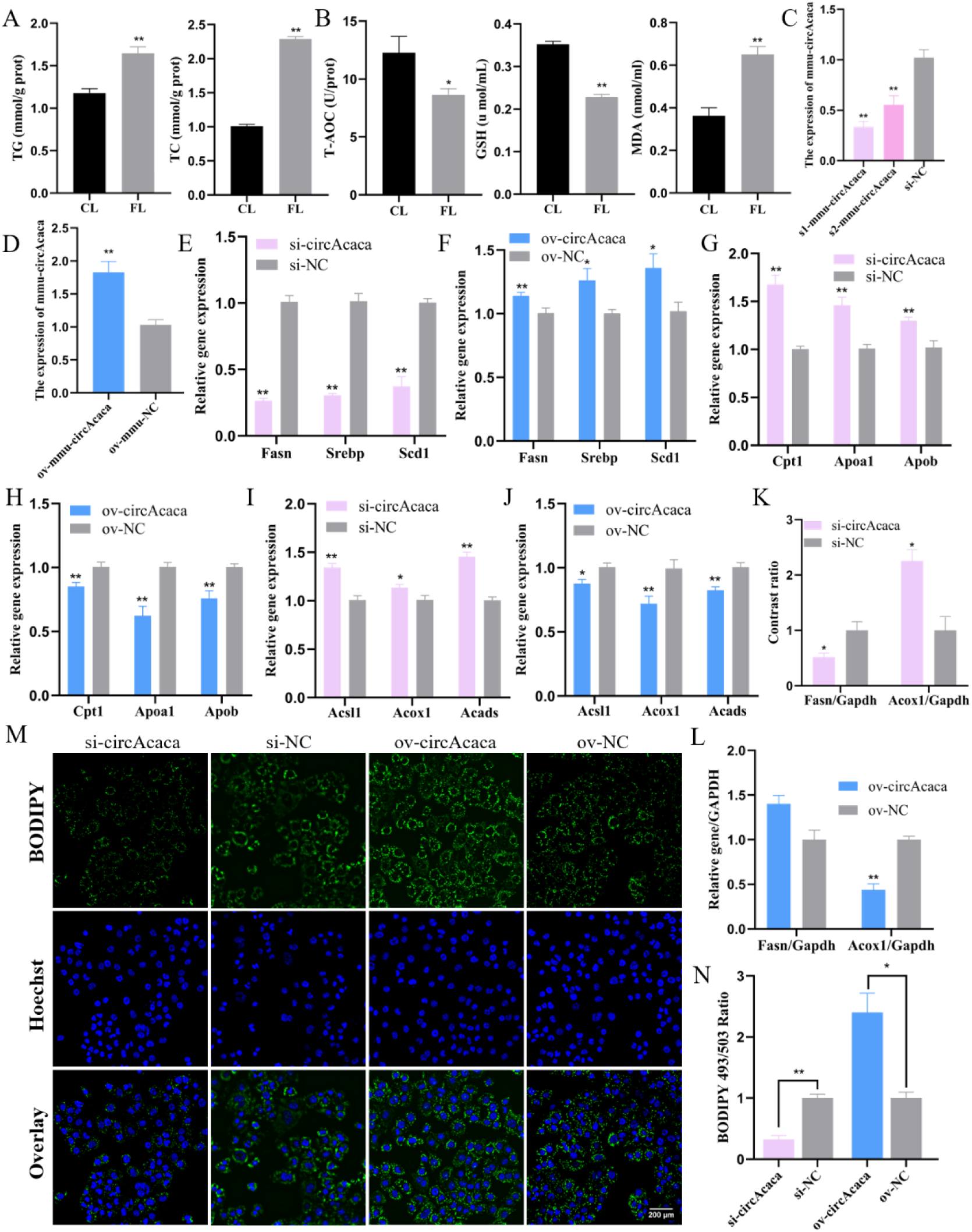
Validation of the functional conservation of mmu-circAcaca in mouse liver cells. (A) Assessment of the content of TG and TC in FL and CL mice serum. (B) Analysis of concentration of T-AOC, GSH and MDA in FL and CL mice serum. (C) qPCR evaluation of the interference efficiency of s1-mmu-circAcaca and s2-mmu-circAcaca on mmu-circAcaca. (D) qPCR analysis of the expression content of mmu-circAcaca after transfection with ov- mmu-circAcaca. (E, F) qPCR measurement of the mRNA expression of Fasn, Srebp and Scd1 after knockdown and overexpression of mmu-circAcaca. (G, H) qPCR assessment of the mRNA content of Cpt1, Apoa1 and Apob after transfection with si-mmu-circAcaca and ov- mmu-circAcaca. (I, J) qPCR analysis of the expression of Acsl1, Acox1 and Acads after interference and upregulated of mmu-circAcaca. (K) Statistics of Fasn and Acox1 protein gray value after knockdown of mmu-circAcaca. (L) BODIPY 493/503 imaging of lipid droplets following the silencing and overexpression of mmu-circAcaca. Scale bars= 200 μ m. (M) Statistics of Fasn and Acox1 protein gray value after overexpression of mmu-circAcaca. (N) Quantification of the BOD 493/503-positive staining area following the alteration of ma expression. The data is represented as mean ± SEM. * *P* < 0.05; ** *P* < 0.01.

**S11 Fig.**
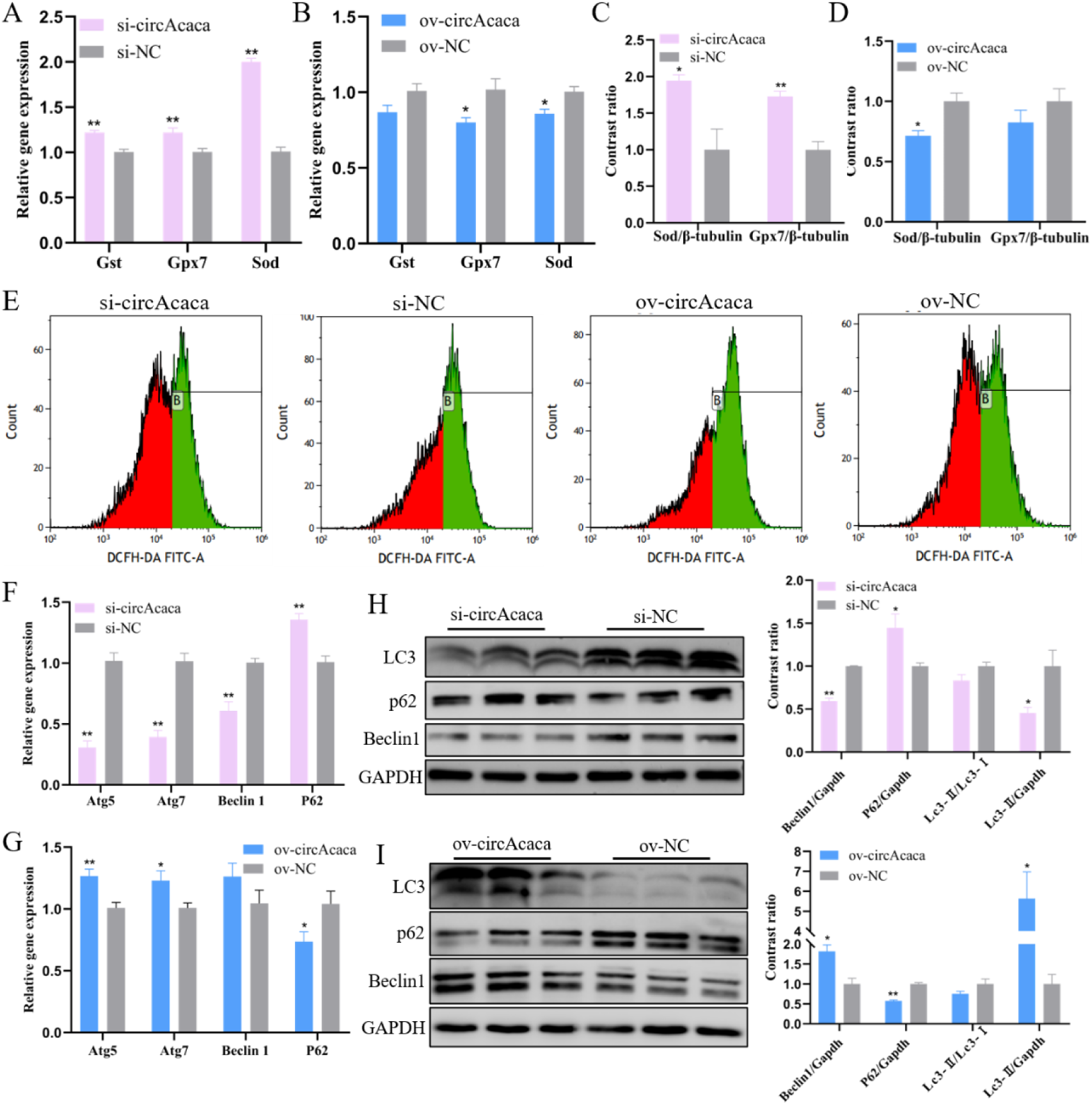
Validation of the functional conservation of mmu-circAcaca in mouse liver cells. (A, B) qPCR assessment of the mRNA content of Gst, Gpx7 and Sod after knockdown and overexpression of mmu-circAcaca. (C, D) Quantification of the protein content of Sod and Gpx7 after changing the expression of mmu-circAcaca. (E) Detection of ROS levels inside cells after the interference and overexpression of mmu-circAcaca. (F, G) qPCR evaluation of the mRNA level of autophagy related genes after silencing and upregulated of mmu-circAcaca. (H, I) Western blot analysis of the protein expression of autophagy related genes after transfection si-mmu-circAcaca and ov-mmu-circAcaca. The data is represented as mean ± SEM. * *P* < 0.05; ** *P* < 0.01.

S1 Table. RNA oligonucleotides in this study. S2 Table. Primers for qPCR.

S3 Table. Antibodies for western blot.

S4 Table. FISH probe sequences in this study.

S5 Table. Summary of data generated from circRNAs deep sequencing.

S6 Table. Different type circRNAs in each sample.

S7 Table. All circRNAs detected in chicken liver (TPM).

S8 Table. All the differentially expressed circRNAs.

S9 Table. Target miRNAs of 8 differentially expressed circRNAs.

S10 Table. The target genes of miR-132b-5p and miR-101-2-5p were predicted by different software.

S11 Table. Conservation analysis of circACACA.

